# Xeno-Free Peptide-Functionalized Hydrogels Support hiPSC Encapsulation and *In Situ* Differentiation into Cardiomyocytes with Enhanced Structural Maturation

**DOI:** 10.64898/2026.07.08.737331

**Authors:** Mohammadjafar Hashemi, Nongmaithem Debeni Devi, Yasaman Kargar Gaz Kooh, Charlotte Chen, Ganesh Malayath, Julia Victor, Bahareh Bahmani, Nathaniel Huebsch

## Abstract

While defined synthetic substrates can replace Matrigel for human induced pluripotent stem cell (hiPSC) culture and hiPSC–derived cardiomyocyte (hiPSC-CM) production, existing approaches culture cells on two-dimensional surfaces and yield structurally immature cardiomyocytes, limiting their use in disease modeling and regenerative medicine. Here, we developed a xeno-free, fully-defined cyclic RGD (cRGD)-functionalized alginate platform in which we encapsulated hiPSCs to support their expansion and *in situ* cardiac differentiation. cRGD functionalization was essential for hiPSC survival and pluripotency, with maximal support achieved at a low ligand density (25 µM). In the presence of cRGD, hiPSC encapsulation into softer gels made from lower molecular weight alginates led to enhanced hiPSC expansion and improved cardiogenesis. Strikingly, differentiation *in situ* with 3D gels led to hiPSC-CM with higher structural maturity, including a markedly increased proportion of cardiomyocytes positive for the structural maturation markers Desmin and MYH7. Finally, after enzymatic retrieval from hydrogels, cardiomyocytes derived from softer gels formed tissue-engineered myocardium with superior contractile stress compared to tissue fashioned from hiPSC-CM derived from more rigid gels. Together, these results demonstrate the promise of this defined, tunable platform for biomanufacturing of structurally more mature cardiomyocytes from hiPSC.

## Introduction

Scalable production of structurally mature human cardiomyocytes from induced pluripotent stem cells (hiPSCs) remains one of the most pressing unmet challenges in cardiac biology, limiting progress in disease modeling, drug screening, and cell-based therapies [1]. Small-molecule based differentiation strategies such as the “GiWi” protocol that relies on Wnt-pathway modulation have standardized hiPSC-to-cardiomyocyte differentiation in two-dimensional (2D) monolayer culture across many laboratories [2, 3]. Yet significant challenges persist: batch-to-batch variability, inconsistent differentiation efficiency, and the inherently limited scalability of monolayer systems continue to constrain the reliable production of high-quality cardiomyocytes at quantities relevant for downstream applications [4–6]. Critically, hiPSC-CMs derived in conventional 2D systems often fail to exhibit proper cellular localization of key structural proteins such as Desmin [7, 8], reflecting the broader problem of structural immaturity that limits their utility for disease modeling and tissue engineering applications [1, 9]. Differentiation in 3D formats such as embryoid bodies (EBs) has been shown to improve both hiPSC-CM production scalability and developmental maturation compared to 2D monolayer culture [10, 11], and more recently, hiPSC-CM derivation directly within 3D hydrogel matrices has been shown to further improve structural maturation and diminish user- and batch-dependent variability beyond EB-based differentiation [12].

A central insight driving the design of 3D hydrogel culture systems for stem cell biology is that the extracellular matrix (ECM) is an active regulator of cell fate [9, 13]. While EB-based differentiation improves upon 2D monolayer culture by providing a 3D cellular microenvironment, this process does not allow researchers to tune the biochemical and mechanical properties of the cellular niche, leaving key regulators of stem cell fate essentially uncontrolled [12]. In contrast, engineered hydrogel matrices offer precise, independent control over the biophysical and biochemical cues that collectively govern lineage commitment [14]. ECM stiffness has been shown to direct cellular lineage commitment [15], with soft matrices favoring cardiomyogenic differentiation over stiffer substrates [16]. Alongside mechanical cues, adhesive ligands that engage integrins are equally important for coordinating cell survival, proliferation, and differentiation, meaning both matrix mechanics and integrin-mediated adhesion must be considered for stem cell differentiation. Peptide motifs such as Arg-Gly-Asp (RGD) serve as such adhesive ligands, activating downstream mechanotransduction and potentiating growth factor signaling pathways [17].

One challenge with using hydrogels for stem cell differentiation is the potential introduction of xenogeneic (animal-derived) materials such as bovine fibrinogen [12]. This is likely to complicate efforts to translate gel-differentiated cells for clinical applications that require transplantation [18]. Thus, biocompatible synthetic or semi-synthetic gel-forming materials are especially attractive. Within this category of materials, alginate hydrogels have long been recognized as a versatile platform for stem cell encapsulation, offering excellent biocompatibility, tunable mechanical properties, and the ability to shield encapsulated cells from mechanical damage in suspension culture [19]. Early work demonstrated that human embryonic stem cells (hESCs) encapsulated in calcium alginate hydrogels could be maintained in an undifferentiated state for extended periods without feeder cells or passaging, while retaining pluripotency [20]. Beyond pluripotency preservation, alginate encapsulation has been shown to actively enhance directed differentiation: Richardson *et al.* demonstrated that encapsulation of undifferentiated hESCs followed by *in situ* differentiation induction resulted in higher viability and a dramatically stronger differentiation response compared to parallel 2D cultures, with alginate encapsulation yielding a 20-fold increase in differentiation efficiency toward pancreatic islet-like cells [21]. These observations suggest that the 3D encapsulation environment itself can potentiate lineage commitment beyond what is achievable in monolayer, motivating its application to cardiac differentiation. However, alginate is inherently bioinert, lacking native cell-adhesion motifs, which limits direct cell-matrix interactions critical for stem cell fate regulation [22]. This motivates functionalization with bioactive peptides such as RGD, which engage integrins to prevent anoikis and activate mechanotransduction signaling [23], alongside modulation of gel stiffness through approaches including control over alginate molecular weight [24] to better recapitulate the mechanical behavior of native cardiac ECM within the developing heart tube.

Here, we explored the possibility of using fully-defined, xeno-free peptide-modified alginate gels for hiPSC-CM biomanufacturing. Cyclic RGD (cRGD) functionalized alginate hydrogels dramatically improved hiPSC expansion while maintaining pluripotency. Importantly, differentiation within gels via the GiWi protocol led to high differentiation efficiencies but with nearly 80% lower batch-to-batch variability in cardiomyocyte purity (coefficient of variation across N = 4 matched differentiation batches) compared to matched monolayer differentiations in an hiPSC line for which monolayer has been optimized in our laboratory for several years [25–27]. Strikingly, differentiation within gels yielded hiPSC-CMs that exhibited hallmarks of structural maturation, including robust desmin expression. We refer to these gel-encapsulated, differentiated constructs as Engineered iPSC Cardiobodies (EiCs). Finally, we demonstrate the gel-derived hiPSC-CM can be used for downstream tissue engineering applications such as formation of micro-engineered heart tissues (µHTs). Together, these findings establish a xeno-free, tunable alginate encapsulation platform that bridges the critical gap between scalable hiPSC biomanufacturing and the production of functionally competent cardiomyocytes for tissue engineering, disease modeling, and drug screening applications.

## Results

### Alginate hydrogels as matrices for hiPSC encapsulation and cardiac differentiation

Hydrogels were formulated by combining cRGD-conjugated alginate (**Fig. 1A**) at varying peptide concentrations (0, 25, and 75 µM) with calcium crosslinker across three alginate molecular weights (60, 135, and 290 kDa), enabling independent tuning of biochemical and mechanical properties of the encapsulation matrix; detailed gel formulation parameters are presented in **Table 1** and **Fig. 1B**. hiPSCs were encapsulated within these hydrogels at day -3 (**Fig. 1C and Fig. S1**). To determine whether calcium used for crosslinking might be detrimental to iPSC viability, we next varied gel crosslinking and media calcium concentrations and performed Calcein-AM/Ethidium Homodimer (live/dead staining). hiPSC viability was comparable at baseline and lower calcium levels, declining only at supra-baseline concentrations (doubling the concentration used for crosslinking, or increasing the amount of calcium in the basal media by a factor of 10; **Fig. S2B, C**). These findings confirm that the calcium concentrations used throughout this study were within an optimal range to maintain hiPSC survival.

**Figure 1.**
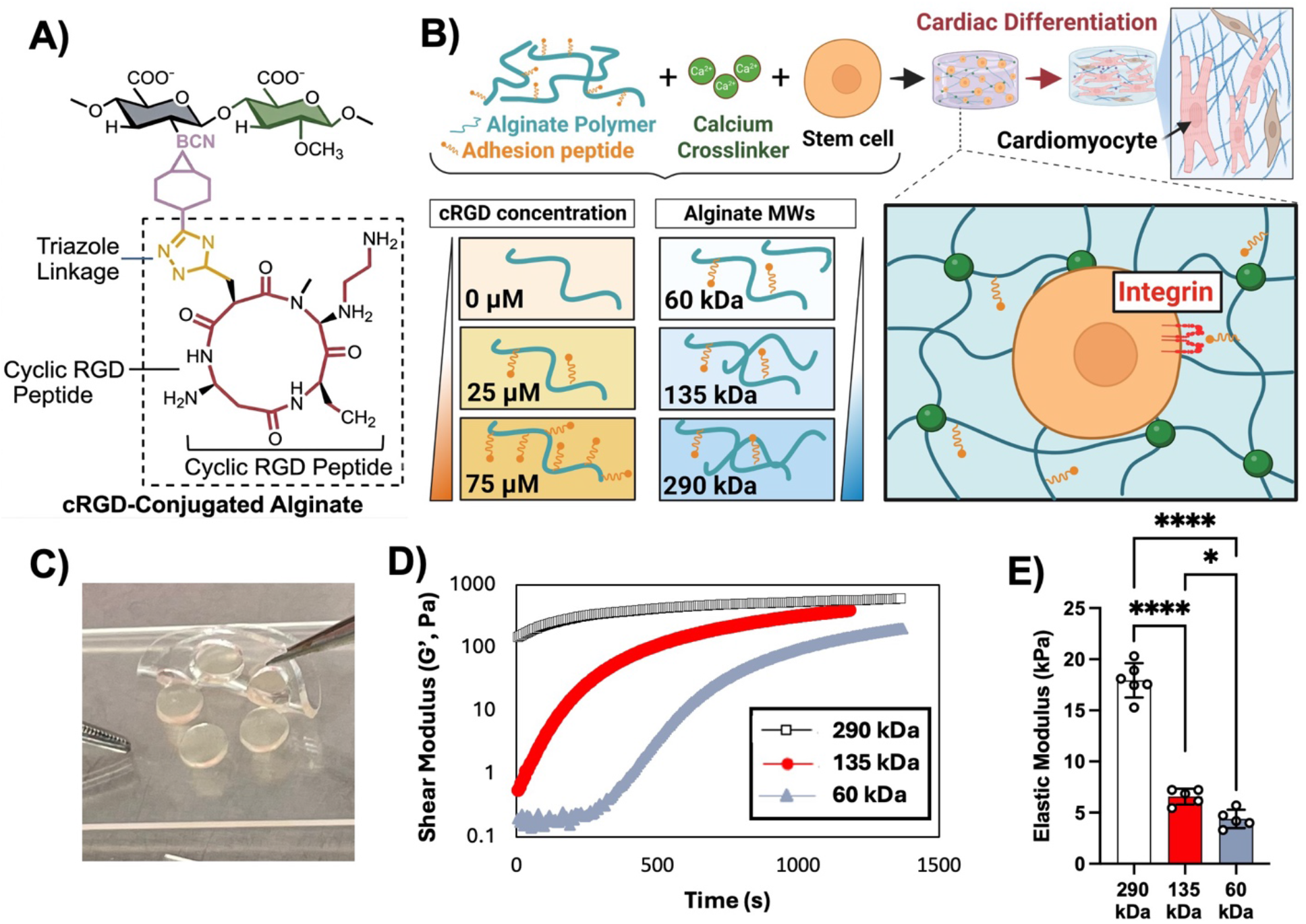
Alginate hydrogels as a matrix for hiPSC encapsulation and cardiac differentiation. **A)** Schematic depicting the SPAAC bioconjugation reaction between azide-terminated cRGD and BCN-modified alginate. **B)** Schematic depicting encapsulation of hiPSCs within alginate hydrogels formulated with varying peptide concentrations and stress-relaxation properties (tuned via alginate molecular weight). **C)** Representative photograph of alginate hydrogel discs formed for hiPSC encapsulation. **D)** Time-dependent evolution of storage modulus (G′) during calcium-mediated gelation of alginate hydrogels with different molecular weights at the same calcium concentration. **E)** Elastic modulus of hydrogels made from alginate with varying molecular weights. Each replicate represents one hydrogel; data are representative of two independently formed hydrogel batches. *p < 0.05, ****p < 0.0001 (one-way ANOVA followed by post hoc Tukey test). Schematic in panel B created with BioRender.

**Table 1:** Alginate Hydrogel Formulation Used for Cell Encapsulation.

| Condition | Naked Alginate (MW) | cRGD-Modified Alginate (MW) | Final cRGD Concentration | Final Alginate Concentration |
|---|---|---|---|---|
| 290 kDa | 290 kDa | — | 0 µM | 1% w/v |
| 290 kDa + 25 µM cRGD | 290 kDa | 290 kDa (150 µM stock) | 25 µM | 1% w/v |
| 290 kDa + 75 µM cRGD | 290 kDa | 290 kDa (150 µM stock) | 75 µM | 1% w/v |
| 135 kDa + 25 µM cRGD | 135 kDa | 290 kDa (150 µM stock) | 25 µM | 1% w/v |
| 60 kDa + 25 µM cRGD | 60 kDa | 290 kDa (150 µM stock) | 25 µM | 1% w/v |

Rheological characterization confirmed that alginate molecular weight influences the rate of network formation and the resulting gel mechanics (**Fig. 1D-E**). Elastic modulus measurements revealed that all three gel formulations were significantly different from one another, with 290 kDa gels exhibiting the highest stiffness (17.9 ± 1.7 kPa), followed by 135 kDa (6.6 ± 0.8 kPa) and 60 kDa gels (4.4 ± 0.9 kPa) (**Fig. 1E**), confirming that each molecular weight formulation produces a mechanically distinct hydrogel environment for hiPSC encapsulation.

### Low-dose cRGD promote hiPSC growth and maintenance of stemness

Brightfield images of hiPSCs encapsulated into 290 kDa alginate at different concentrations of cRGD revealed differences in colony morphology across cRGD concentrations, with more defined colony boundaries and lumen-like structures visible at 25 µM cRGD after 3 days of expansion (**Fig. 2B**) and 5 days of expansion (**Fig. S2A**). hiPSCs encapsulated without cRGD (0 µM) exhibited notably lower viability compared to hiPSC in both peptide-functionalized gel types (**Fig. 2C**). Premature differentiation (typically down an ectodermal lineage [28]) can limit iPSCs’ ability to develop into cardiomyocytes. Thus, we assessed pluripotency via immunofluorescence analysis of Nanog, one of the first transcriptional markers of iPSC to be deactivated during differentiation [29]. This analysis revealed that pluripotency was markedly reduced in gels without cRGD modification after only 3 days of culture (0 µM condition; **Fig. 2D**). In contrast, robust Nanog-positive staining was observed throughout the gel in both 25 µM and 75 µM cRGD conditions, confirming that cRGD functionalization is necessary for maintaining hiPSC pluripotency within alginate matrices.

**Figure 2.**
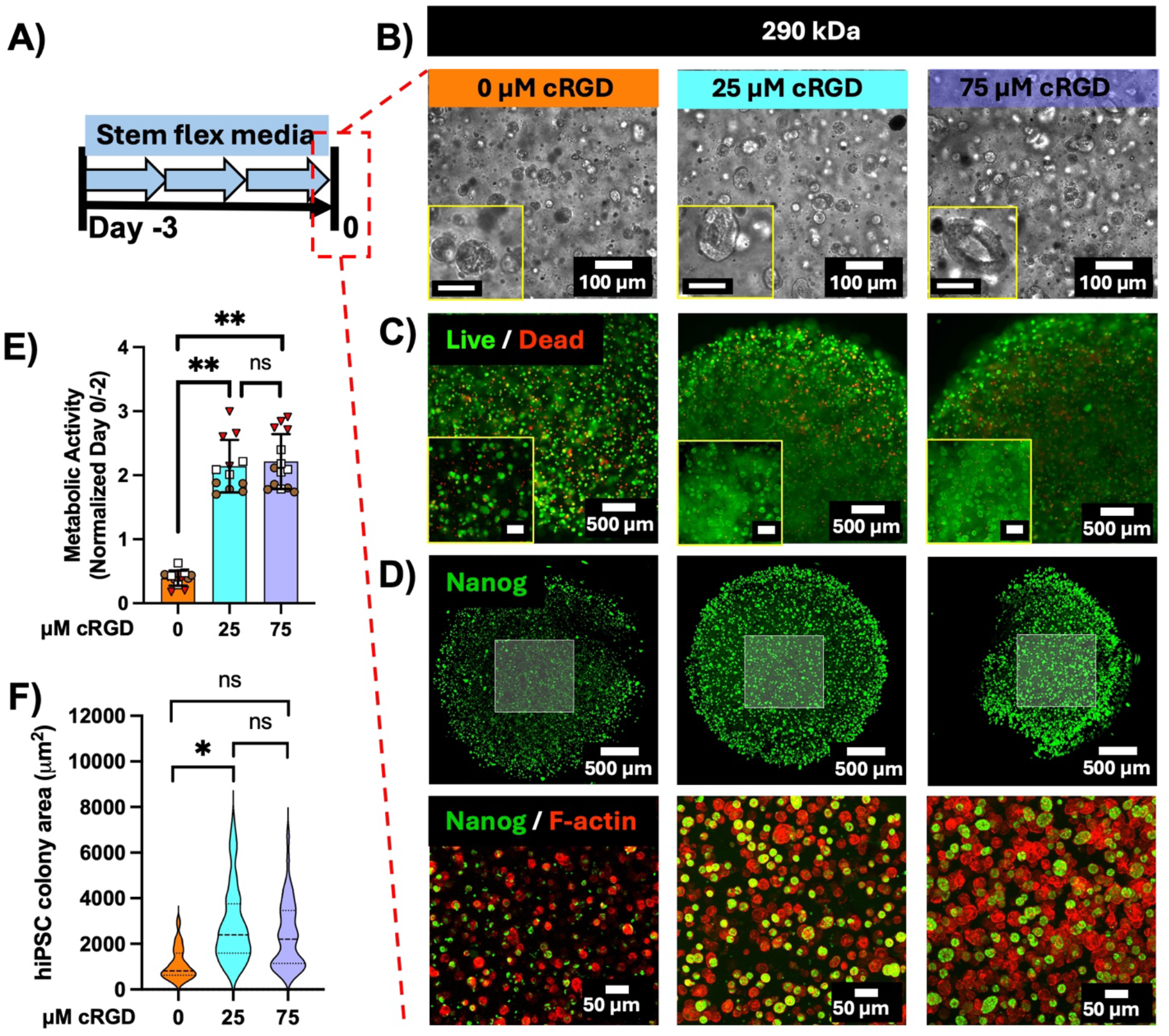
Low-dose cRGD promotes hiPSC growth and maintenance of stemness. **A)** Timeline of culturing hiPSC-laden alginate for three days in Stem Flex media. Representative **B)** brightfield, **C)** live/dead fluorescence, and **D)** Nanog immunofluorescence images of encapsulated hiPSCs in 290 kDa alginate gels at day 0 (three days post-encapsulation) across cRGD concentrations (0, 25, and 75 µM). Scale bars = 50 µm (brightfield) and 200 µm (live/dead). **E)** Quantification of resazurin reduction at day 0, normalized to day −2, across cRGD concentrations (0, 25, and 75 µM); symbols indicate individual gels grouped by independent encapsulation batch. Data are presented as mean ± SD; N = 3 independent encapsulation batches per condition (n = 3 to 5 gels per batch); nested one-way ANOVA with Tukey’s multiple comparisons test; ns, not significant; **p < 0.01. **F)** Quantification of colony area across cRGD concentrations (0, 25, and 75 µM). Data are presented as mean ± SD; N = 3 independent encapsulation batches per condition (n = 44, 188, and 155 colonies for 0, 25, and 75 µM, respectively); nested one-way ANOVA with Tukey’s multiple comparisons test; *p < 0.05; ns, not significant.

To assess the effect of cRGD concentration on hiPSC metabolic activity following encapsulation, we measured resazurin reduction at day 0 and normalized to day −2 baseline measurements (**Fig. 2E**). Both cRGD concentrations significantly enhanced normalized metabolic activity compared to the no-peptide control (**Fig. 2E**), with no additional benefit observed at 75 µM over 25 µM (*p* > 0.9), suggesting 25 µM is sufficient for maximal metabolic support and/or hiPSC expansion. As resazurin reduction integrates both metabolic activity per cell and overall cell number, we directly assessed hiPSC expansion by quantifying cross-sectional colony area within the gels. Colony area was significantly larger at 25 μM than at 0 μM cRGD, whereas 75 μM did not differ significantly from either condition (**Fig. 2F**), indicating that a low cRGD density is sufficient to promote hiPSC colony growth within the alginate matrix.

### Softer gel supports better hiPSC expansion and self-organization

Morphologically, hiPSCs encapsulated in stiffer 290 kDa gels formed smaller colonies, while cells in softer 60 kDa gels exhibited larger colonies (**Fig. 3A**). hiPSC colony area was tracked across day −2, −1, and 0, becoming significantly larger in 60 kDa compared to 290 kDa gels by day 0 (**Fig. 3B**), with no significant difference between conditions at the earlier timepoints. Lumen-like structures became significantly more prevalent over time in softer gels, with a greater percentage of 60 kDa colonies exhibiting a lumen by day 0 compared to 290 kDa (**Fig. 3C and Fig. S3G**); among lumen-containing colonies, the lumen area to colony area ratio was also significantly higher in 60 kDa gels (**Fig. 3D**), while colony circularity, lumen area, and lumen thickness did not differ significantly across molecular weights (**Fig. S3D–F**), consistent with prior work on lumen formation in alginate-encapsulated hiPSCs [30]. By differentiation day 2, lumen-like structures were no longer apparent in any condition (**Fig. S3H**), suggesting that the lumen space is rapidly filled in by proliferating cells shortly after the onset of differentiation.

**Figure 3.**
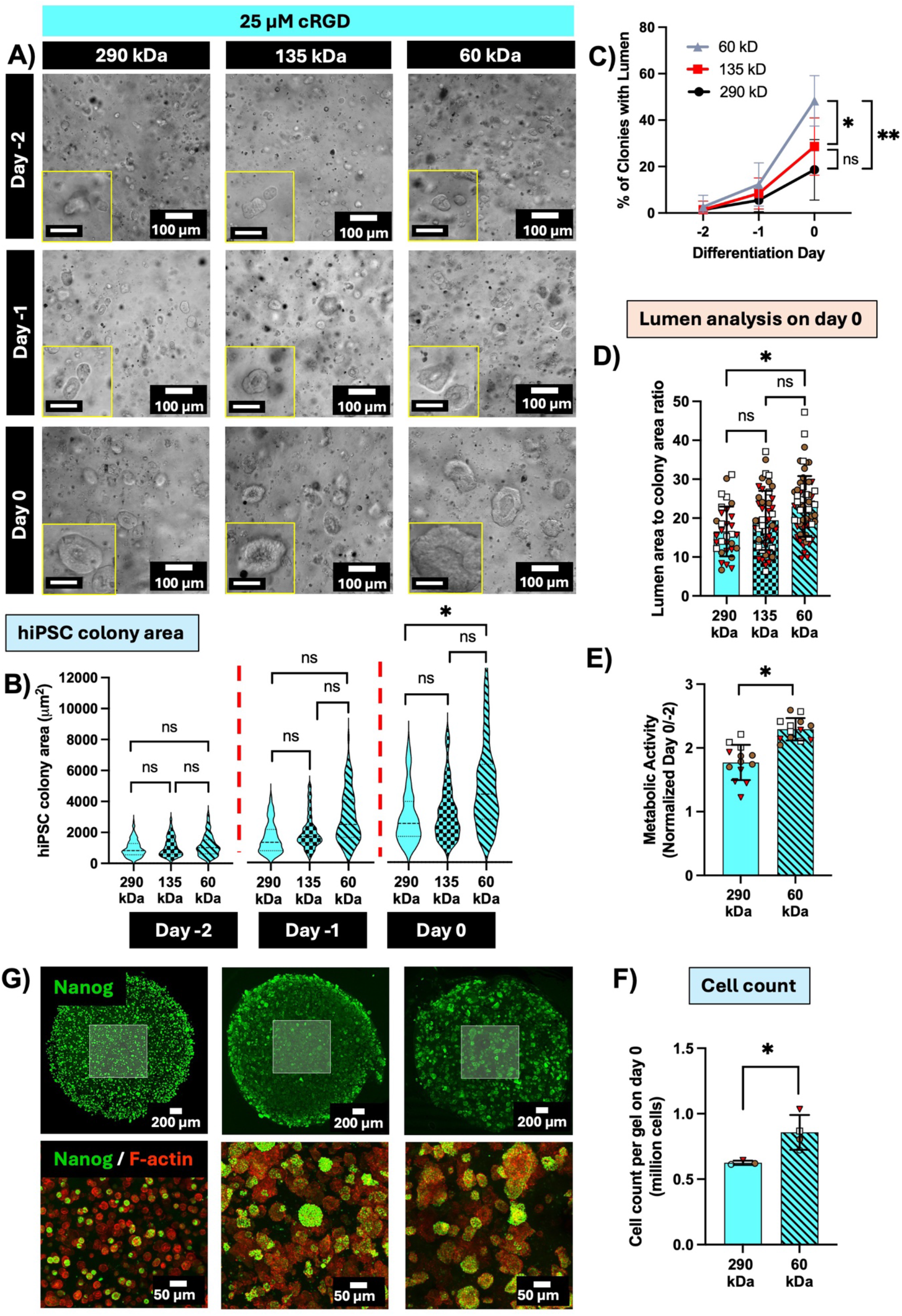
Softer gels support hiPSC expansion and self-organization. **A)** Representative brightfield images of hiPSC-laden alginate hydrogels at day −2, day −1, and day 0 across three alginate molecular weights (290, 135, and 60 kDa) at 25 µM cRGD. Insets show higher magnification views of individual colonies. Scale bars = 50 µm. **B)** hiPSC colony area at day −2, day −1, and day 0; n = 191, 212, and 222 colonies at day −2; n = 213, 180, and 173 at day −1; n = 188, 162, and 130 at day 0, for 290, 135, and 60 kDa, respectively; N = 3 independent encapsulation batches per condition. Nested one-way ANOVA with Tukey’s multiple comparisons: day −2 and day −1, ns for all comparisons; day 0, 290 kDa vs. 60 kDa, *p = 0.0331, all other day 0 comparisons ns. **C)** Percentage of colonies with a lumen-like structure across conditions and days; n = 23, 23, and 21 gels for 290, 135, and 60 kDa; N = 3 independent encapsulation batches per condition.* p = 0.0402, ** p = 0.0068, Nested 1-way ANOVA followed by Tukey’s multiple comparison test, comparing different polymer conditions at day 0), **D)** Lumen area to colony area ratio among lumen-containing colonies at day 0; each dot represents one colony (n = 34, 47, and 58 colonies; N = 3 independent encapsulation batches per condition). **E)** Metabolic activity in 290 kDa and 60 kDa gels at day 0, normalized to day −2; each dot represents one gel (n = 13 gels per condition; N = 3 independent encapsulation batches per condition). Nested t test, *p = 0.0449. **F)** Cell count per gel at day 0 in 290 kDa and 60 kDa gels; each dot represents one independent encapsulation batch (N = 3, 290 kDa; N = 4, 60 kDa). Unpaired t test with Welch’s correction, *p = 0.0382. **G)** Nanog immunofluorescence (top) and Nanog/F-actin merge (bottom) images at day 0. Nested t test was used for B–F; ns, not significant; * p < 0.05; ** p < 0.01.

Just as increasing peptide density increased hiPSC number and/or per-cell metabolic activity (**Fig. 2**), decreasing polymer M_w_ also increased resazurin reduction (**Fig. 3E**). Metabolic activity was significantly higher in hiPSC within 60 kDa gels as compared to hiPSC in 290 kDa gels (nested *t*-test, *p = 0.0449). Increased resazurin reduction for hiPSC in the 60 kDa gels may reflect an ability of the cells to mechanically reorganize these softer matrices [24], consistent with the greater colony spreading observed in 60 kDa gels described above. Cell count per gel at day 0 was also significantly higher in 60 kDa compared to 290 kDa gels (**Fig. 3F**), providing a direct, cell-number-based confirmation that softer gels support greater hiPSC expansion. Notably, although 60 kDa + 25 µM cRGD-modified gels out-performed 290kDa + 25 µM cRGD-modified gels in supporting hiPSC expansion, 290 kDa + 25 µM cRGD-modified gels still out-performed lower M_w_ gels lacking cell-adhesive peptides, demonstrating a critical role for integrin-mediated cell-material interaction in hiPSC expansion and indicating that peptide presentation influenced early expansion more strongly than alginate molecular weight (**Fig. S3 A, B**). Even without cell adhesive peptides, however, 60 kDa gels still out-performed embryoid bodies in terms of their ability to promote hiPSC expansion (**Fig. S3A**), potentially by protecting the growing hiPSC colonies from shear-induced damage during routine media changes. Nanog-positive colonies were observed across all three molecular weight conditions, confirming retention of pluripotency marker expression within the alginate matrix regardless of gel stiffness (**Fig. 3G**).

### Differentiation in Soft hydrogels enhance hiPSC-cardiomyocyte production

Having demonstrated an ability to expand iPSC whilst maintaining pluripotency within cRGD-modified alginates, we next investigated the possibility of using these materials for hiPSC-CM biomanufacturing. Following initial expansion in stem cell media, we initiated cardiomyocyte differentiation (**Fig. 4A**) using the standard “GiWi” protocol [2, 31]. Similar to prior work [12, 32, 33], we observed that as the iPSC differentiated, they tended to form contiguous “tissue-like” structures within the surrounding matrix (**Fig. 4I**). To directly assess contractile function within the intact hydrogel, we measured peak contraction velocity of beating EiCs *in situ*, finding that EiCs derived from softer 60 kDa gels contracted significantly faster than those derived from 290 kDa gels (*p < 0.05; **Fig. 4B and Fig. S4C**), indicating that the contractility benefits of softer gels are already apparent prior to gel dissociation and µHT formation.

**Figure 4.**
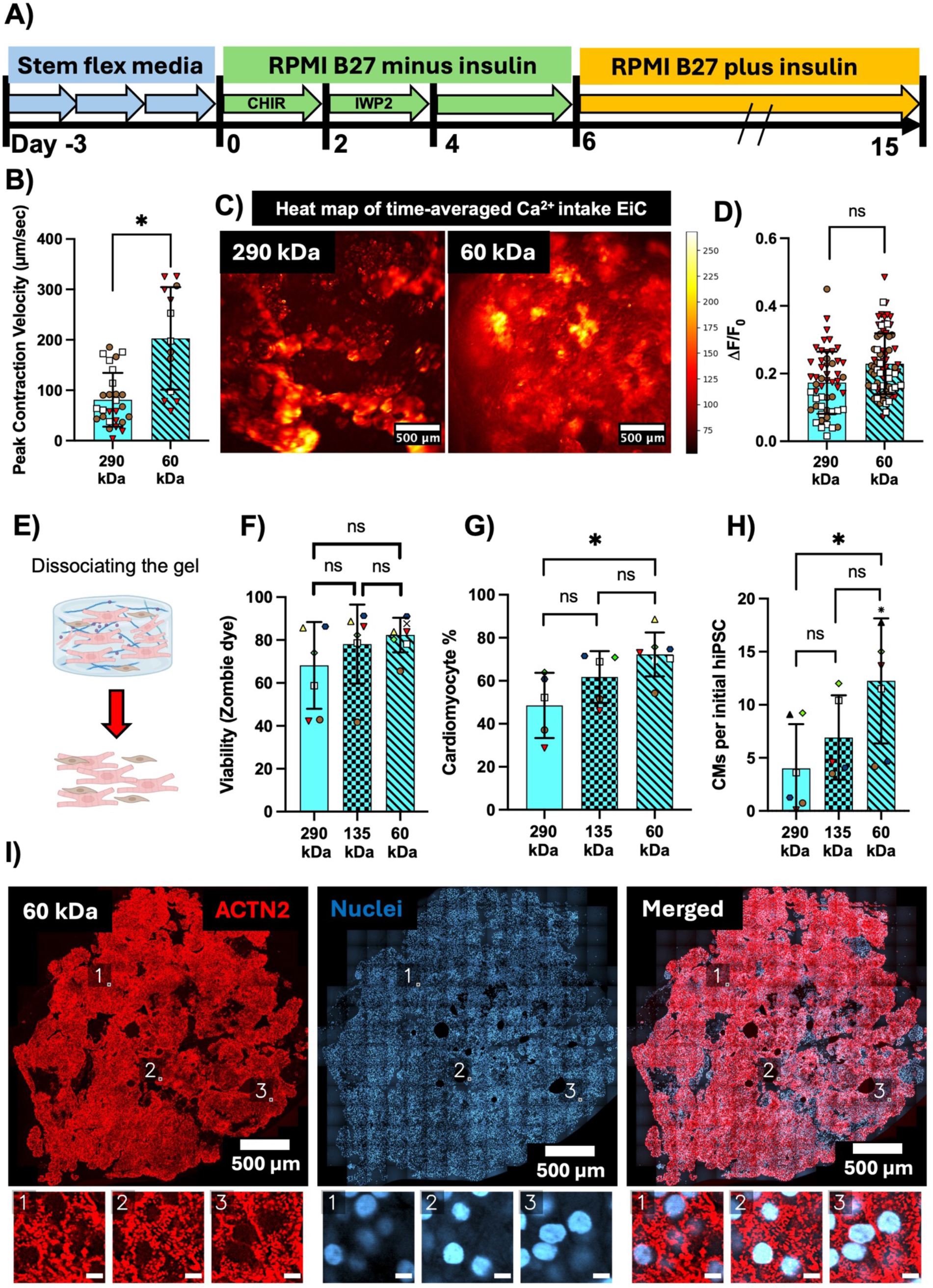
Softer hydrogel enhances cardiomyocyte production. **A)** Cardiac differentiation timeline: Stem Flex media (day −3 to 0), RPMI B27 minus insulin with CHIR (day 0–2) and IWP2 (day 2–4), and RPMI B27 plus insulin (day 6–15). **B)** Peak contraction velocity of EiCs derived from 290 kDa and 60 kDa alginate gels at 25 µM cRGD; each dot represents one gel (n ≥ 13 gels per condition, pooled from N = 3 independent encapsulation and differentiation batches). **C)** Representative heatmaps of time-averaged Ca²⁺ intake in EiCs derived from 290 kDa and 60 kDa alginate gels at 25 µM cRGD. **D)** Calcium transient amplitude (ΔF/F0) in EiCs derived from 290 kDa and 60 kDa alginate gels; each dot represents one ROI recording ( = 53 ROIs, 290 kDa; n = 74 ROIs, 60 kDa; N = 3 independent encapsulation batches per condition). **E)** Schematic of gel dissociation for retrieval of hiPSC-derived cardiomyocytes. **F)** Cell viability following gel dissociation across three alginate molecular weights, assessed by Zombie dye staining. **G)** Cardiomyocyte purity (%) and **H)** yield (CMs per initial hiPSC seeded) across three alginate molecular weights. **I)** Representative immunofluorescence images of EiCs derived from 60 kDa alginate gels stained for ACTN2 (red) and nuclei (blue), shown separately and merged; numbered insets show higher magnification views of sarcomeres. Scale bars = 500 µm (whole-gel) and 5 µm (insets). Statistical comparisons: F) Kruskal-Wallis test, N = 7, 6, and 8 independent batches for 290, 135, and 60 kDa, respectively (ns); G) Kruskal-Wallis test with Dunn’s multiple comparisons test, N = 7, 7, and 10 independent batches for 290, 135, and 60 kDa, respectively (*p = 0.0105, 290 vs. 60 kDa; all other comparisons ns); H) one-way ANOVA with Tukey’s multiple comparisons test, N = 6, 5, and 7 independent batches for 290, 135, and 60 kDa, respectively (*p = 0.0215, 290 vs. 60 kDa; all other comparisons ns). Schematic in panel E created with BioRender.

We next visualized calcium dynamics within the “Engineered iPSC Cardiobodies” (EiCs), via GCaMP6f fluorescence (**Supplementary videos 1-3**) imaging as a gross metric of overall differentiation efficiency (**Fig. 4C**). The heatmap displays the time-averaged GCaMP fluorescence intensity across the EiC in response to external pacing, where warmer colors (yellow) reflect regions of higher cumulative calcium signal and darker colors (dark red) indicate areas of lower average calcium activity. Given that non-cardiomyocytes do not produce rapidly-cycling calcium transients, areas with low time-averaged GCaMP can be assumed to have a low differentiation efficiency, whereas higher fluorescence would be expected to correspond to greater local hiPSC-CM content. Notably, just as 60 kDa gels supported more hiPSC expansion (**Fig. 3**), these softer gels also led to a more uniform and intense time-averaged GCaMP6f signal compared to 290 kDa gels, suggesting higher and more homogeneous cardiomyocyte yield (**Fig. 4C**). Quantification of background-corrected calcium transient amplitude (ΔF/F₀) between 290 kDa and 60 kDa EiCs, analyzed using a nested framework, did not reveal a statistically significant difference (ns; **Fig. 4D**), despite the trend toward the 60 kDa EiC exhibiting stronger overall calcium influx, both in terms of the magnitude of the signal and the spatial prevalence. Consistent with the likelihood that this calcium signal is proportional to the local density of iPS-CM, when batches were matched for cardiomyocyte content (∼ 60%) obtained with an independent assay (FACs), the trend toward differences in ΔF/F₀ vanished (**Fig. S4B**).

Given that Ca^2+^ imaging indirectly suggested increased cardiogenesis in the softest gels, we next sought to directly assess differentiation efficiency. Thus, we dissociated EiC via collagenase treatment (**Fig. 4E**). Since retrieval of viable cells from the gel is critical for accurate flow cytometric analysis of purity, we assessed cell viability following enzymatic dissociation using Zombie dye exclusion (**Fig. 4F**). This analysis indicated that cell viability was comparable across hiPSC-CM retrieved from gels made from all three-alginate M_w_. Consistent with the more prevalent and extensive Ca²⁺ signal observed in 60 kDa gel-derived EiCs, cardiomyocyte purity assessed by flow cytometry was significantly higher in 60 kDa gels compared to 290 kDa gels (**Fig. 4G**), confirming that softer matrices better support cardiogenesis. As several applications of iPS-CM, including transplantation and drug screening, require massive numbers of cells, CM yield is a critical parameter in biomanufacturing. Thus, we also quantified the number of CMs produced per initially encapsulated hiPSC (**Fig. 4H**). Here too, EiCs derived from 60 kDa alginate gels exhibited a greater yield of cardiomyocytes per initial hiPSC than EiCs derived from the more rigid 290 kDa gels, demonstrating that softer gels can produce more cardiomyocytes from the same number of starting cells for these downstream applications. To report the absolute scale of cardiomyocyte production, we additionally quantified the total number of hiPSC-CMs generated per batch across conditions (**Fig. S4E**). The total CM number was significantly higher in softer gels than in stiffer gels (mean: ∼9.6 million, 290 kDa; ∼13.8 million, 135 kDa; ∼24.5 million, 60 kDa, per batch of 2 × 10⁶ input hiPSC).

Prior studies involving *in situ* differentiation within hydrogels have suggested improved differentiation efficiency in gel shapes with a higher surface-to-volume ratio, perhaps due to limitations of nutrient transport into the gels during differentiation [33]. Although our Ca^2+^ imaging did not suggest spatial heterogeneity in the local intensity of the GCaMP6f signal within 60 kDa gel-derived EiC (**Fig. 4C**), we sought to determine the potential for spatially varied hiPSC-CM differentiation via whole mount staining of the gels. Immunostaining for Sarcomeric α-Actinin (ACTN2), revealed widespread, and relatively homogenous distribution of hiPSC-CM within these gels at differentiation day 15 (**Fig. 4I and Supplementary Video 10**). Higher magnification insets revealed defined sarcomeric structures, confirming successful cardiomyocyte differentiation.

Like others in the field [25, 34–36], we typically use hiPSC-CM differentiated in 2D monolayers to fashion “*in vivo*-like” engineered heart muscle. For this work, we routinely apply “quality control” criteria based in part on hiPSC-CM purity. Consistent with our extensive and longstanding use of the WTC-GCaMP as the isogenic control for these studies [26, 37–39], we observe overall high efficiency of the 2D differentiation (**Fig. S4F**). Notably, however, we observed that without extensive optimization, differentiation in EiC made from 60 kDa alginates achieved a similar hiPSC-CM purity and yield as compared to optimized 2D differentiation – albeit with markedly lower (80% reduction in) batch-to-batch variability in cardiomyocyte purity (coefficient of variation across N = 4 matched differentiation batches) (**Fig. S4F and Table S.3**).

### Comparison of cardiomyocyte structural maturation between the best 2D monolayer differentiation and EiC

Lower-batch-to-batch variability in cardiomyocyte purity and yield may be the primary benefit of differentiation in 3D gels, without an impact on the state of individual hiPSC-CM. Thus, we sought to understand whether differentiation in EiC would yield hiPSC-CM with similar structural maturation as cardiomyocytes derived from *optimal* 2D differentiations (defined as having a purity >60 % cTnT at differentiation day 15). We thus assessed markers of structural maturation hiPSC-CM replated to 2D substrates after enzymatic retrieval either from 1) 2D monolayers or 2) gel-based EiC differentiations. Strikingly, hiPSC-CM derived from EiC formed in 60 kDa gels exhibited a significantly lower proportion of MLC2a-positive cardiomyocytes compared to the 2D monolayer derived controls (**Fig. 5A**). Given that MLC2a is expressed in immature ventricular cardiomyocytes and become predominantly restricted to atrial cardiomyocyte during development [40, 41], its reduced expression in hiPSC-CM derived from 60 kDa EiCs (**Fig. 5B**; p = 0.0196) is consistent with maturation-associated changes compared to conventional 2D monolayer differentiation, although a shift toward ventricular specification cannot be excluded. This observation is also consistent with prior transcriptomic comparison of 2D monolayer to 3D embryoid body (EB)-based differentiation by Branco *et al.* [11].

**Figure 5.**
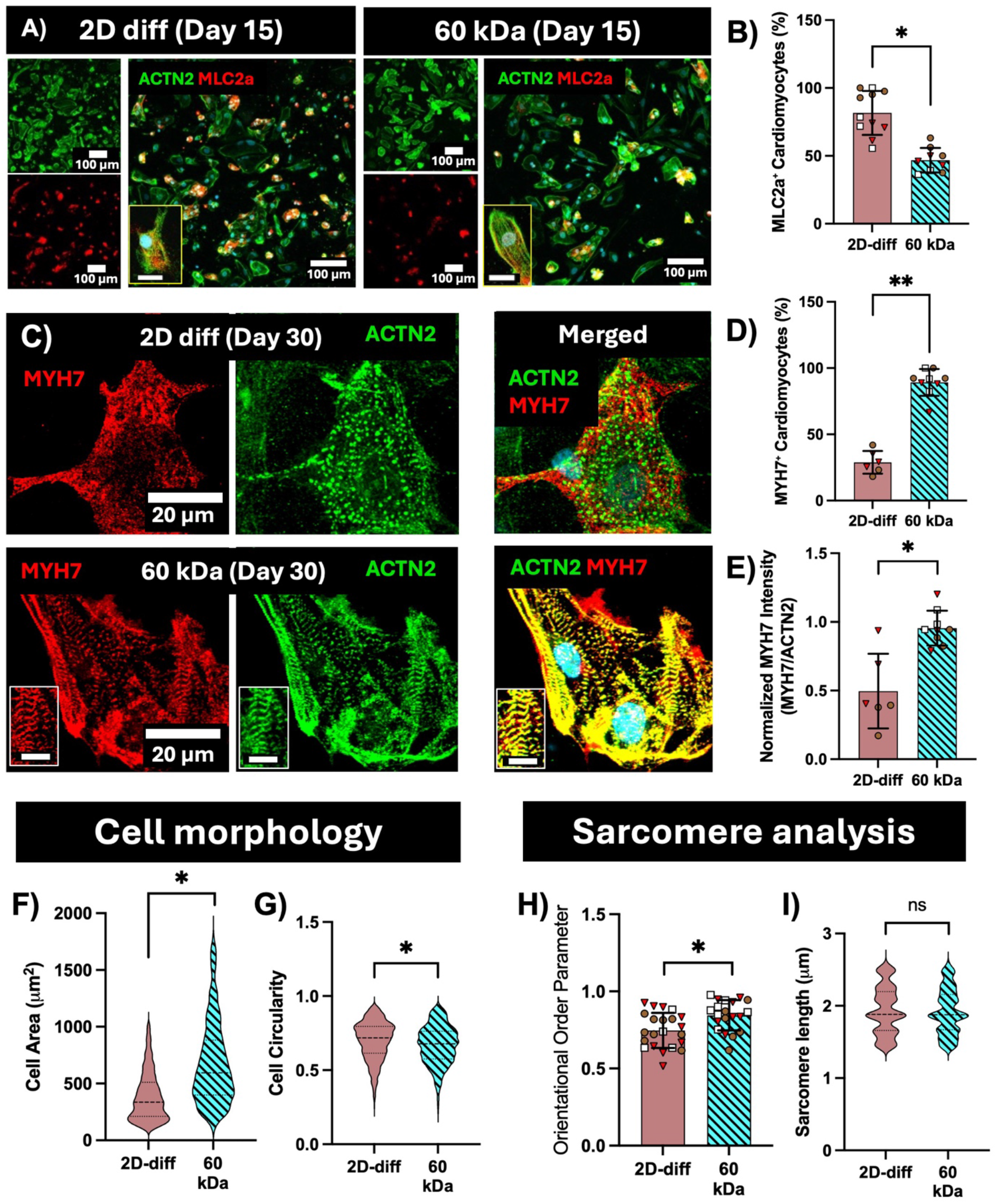
3D encapsulation in soft alginate promotes structural cardiomyocyte maturation. **A)** Representative immunofluorescence images of dissociated cardiomyocytes from 2D monolayer differentiation and 60 kDa EiCs, stained for ACTN2 (green) and MLC2a (red), shown separately and merged. Scale bars = 100 µm, 20 µm (insets). **B)** MLC2a⁺ cardiomyocytes as a percentage of total cardiomyocytes; each dot represents one well, colored by batch (N = 3 independent differentiation batches per condition; n = 11 wells, 2D-diff; n = 9 wells, 60 kDa). **C)** Representative immunofluorescence images of cardiomyocytes stained for MYH7 alone, ACTN2 alone, and merged at day 30; insets show higher magnification views of sarcomeric banding. Scale bars = 20 µm (main images and insets). **D)** MYH7⁺ cardiomyocytes as a percentage of total cardiomyocytes; each dot represents one ROI (N = 2 independent differentiation batches, 2D-diff; N = 3 independent differentiation batches, 60 kDa; n = 15 ROIs total pooled across conditions). **E)** MYH7 intensity normalized to ACTN2 intensity; each dot represents one ROI (N = 2 independent differentiation batches, 2D-diff; N = 3 independent differentiation batches, 60 kDa; n = 15 ROIs total pooled across conditions). **F)** Projected cell area and **G)** cell circularity of replated cells at day 30; each dot represents one cell (N = 3 independent differentiation batches per condition; total n = 2041 cells, F; 2111 cells, G, pooled across conditions). **H)** Orientational order parameter; each dot represents one cell (N = 3 independent differentiation batches per condition; n = 22, 2D-diff; n = 20, 60 kDa). **I)** Sarcomere length; each dot represents one sarcomere (N = 3 independent differentiation batches per condition; total n = 1203 sarcomeres pooled across conditions). Statistical tests: nested t test was used for B, D, E, F, G, and H. Significance: B) *p = 0.0196. D) **p = 0.0015. E) *p = 0.0297. F) *p = 0.0126. G) *p =0.0126. H) *p = 0.0411. I) ns, p =0.8748.

To further assess structural maturation, we stained replated cardiomyocytes for MYH7 (β-myosin heavy chain), a marker of mature, ventricular-like cardiomyocyte phenotype, together with ACTN2 (**Fig. 5C and Fig. S5**). hiPSC-CM derived from EiC formed in 60 kDa gels exhibited a significantly higher proportion of MYH7-positive cardiomyocytes compared to the 2D monolayer derived controls (nested t test, p = 0.0015; **Fig. 5D**). Positive Quantification of MYH7 intensity normalized to ACTN2 intensity revealed significantly higher relative MYH7 expression in hiPSC-CM derived from 60 kDa EiCs compared to 2D monolayer controls (nested t test, p = 0.0491; **Fig. 5E**), further supporting enhanced structural maturation in EiC-derived cardiomyocytes.

Besides lower MLC2a expression and higher MYH7 content, hiPSC-CMs retrieved from 60 kDa EiCs exhibited significantly larger cell area, lower cell circularity, along with higher orientational order parameter compared to 2D-derived cells (**Figs. 5F, 5G, 5H**), indicative of a more rod-like morphology, consistent with a more mature cardiomyocyte structural phenotype. Sarcomere length did not differ significantly between 60 kDa EiC- and 2D-derived cardiomyocytes (**Fig. 5I**), while the orientational order parameter, a measure of sarcomeric alignment, was significantly higher in 60 kDa EiC-derived cardiomyocytes (**Fig. 5H**), indicating more organized sarcomeric structure despite comparable sarcomere length. Together, these findings support the conclusion that 3D encapsulation in 60 kDa alginate promotes a more structurally mature cardiomyocyte phenotype compared to what is achieved in conventional 2D differentiation.

### hiPSC-CMs derived from soft gels exhibit improved functional output in engineered tissues

To evaluate the functional potential of hiPSC-CMs produced in EiCs for downstream applications, dissociated cardiomyocytes from EiCs were seeded to form Micro Heart Tissue (µHT) [25] at day 15 of differentiation, and subsequently cultured until day 26 (**Fig. 6A and S6C**). Importantly, the ability to successfully retrieve viable cardiomyocytes from all three EiC conditions and form functional µHTs demonstrates the feasibility of using EiC-derived hiPSC-CMs for downstream tissue engineering applications. For benchmarking purposes, hiPSC-CMs from the three best-performing 2D monolayer differentiation batches (cTnT > 60 %; **Table 2**) were used as a reference, representing an optimized standard that is difficult to consistently achieve **(Fig. 5 B and C**).

**Figure 6.**
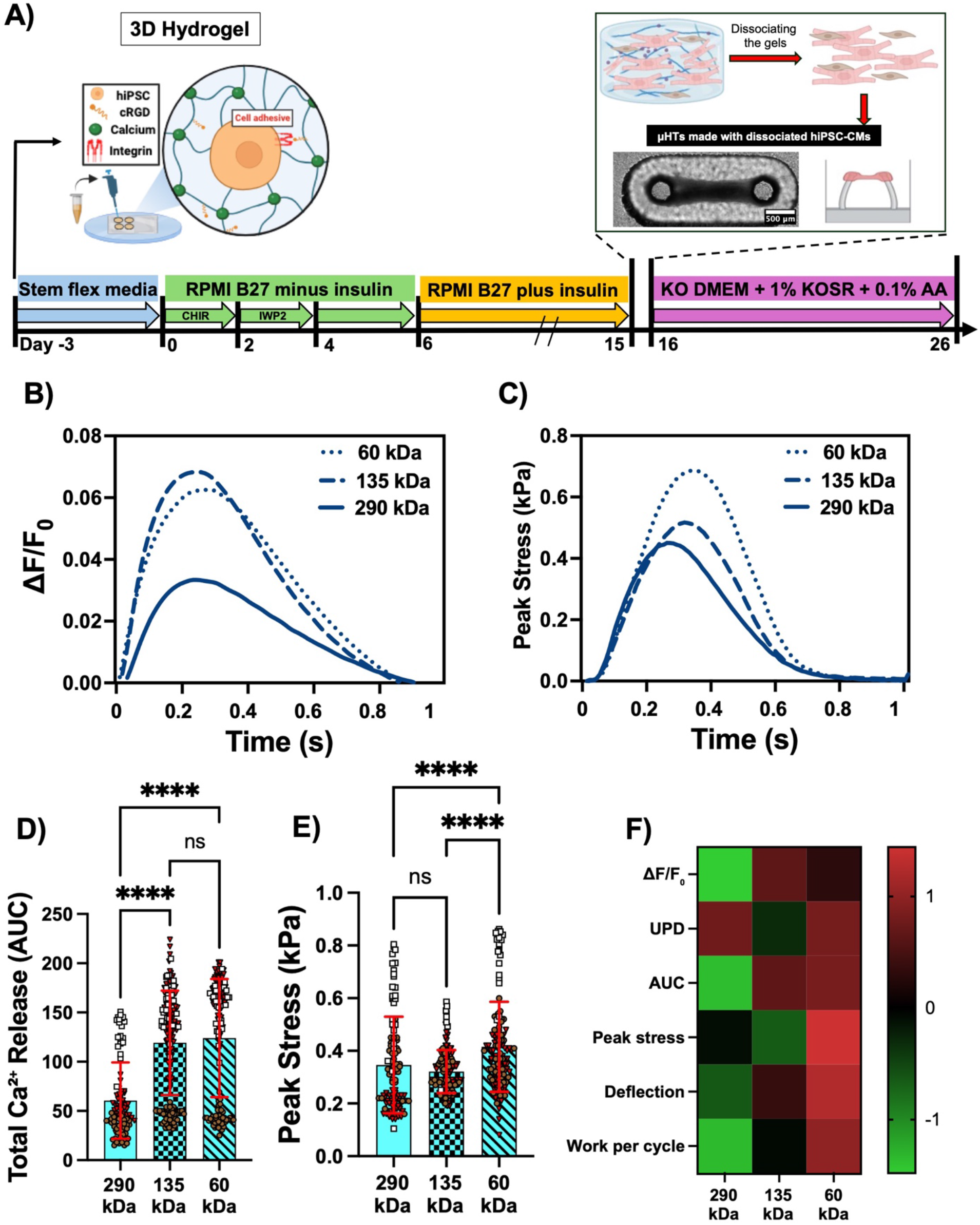
Softer gel-derived hiPSC-CMs improve the functional output of EiC-based micro heart tissues (µHTs). **A)** Schematic of µHT fabrication timeline: 3D hydrogel encapsulation differentiation, gel dissociation, cardiomyocyte retrieval, and µHT formation (day 15–26) in KO DMEM supplemented with 1% KOSR and 0.1% AA. Scale bar = 500 µm. **B)** Representative GCaMP calcium transient traces and **C)** peak stress traces from µHTs across the three conditions. **D)** Total Ca²⁺ release (AUC) and **E)** peak stress (kPa) in µHTs fabricated from cardiomyocytes derived from three alginate molecular weights (290, 135, and 60 kDa); each dot represents one µHT. Because each µHT contains a large number of cells and is formed independent of other µHT, even from the same input batch of cardiomyocytes, each tissue was treated as a separate replicate. µHT data are pooled from *n*=3 independent differentiation batches, and different symbols denote the batch that each µHT came from. Kruskal-Wallis test with Dunn’s multiple comparisons, **** p < 0.0001. **F)** Heatmap of normalized functional metrics across the three conditions (ΔF/F₀, UPD, AUC, peak active force, deflection, and work per cycle; values normalized to −1 to 1). Different letters indicate statistically significant differences between groups (p < 0.05; ANOVA followed by Dunn’s post-hoc test). Schematic in panel A created with BioRender.

**Table 2:**
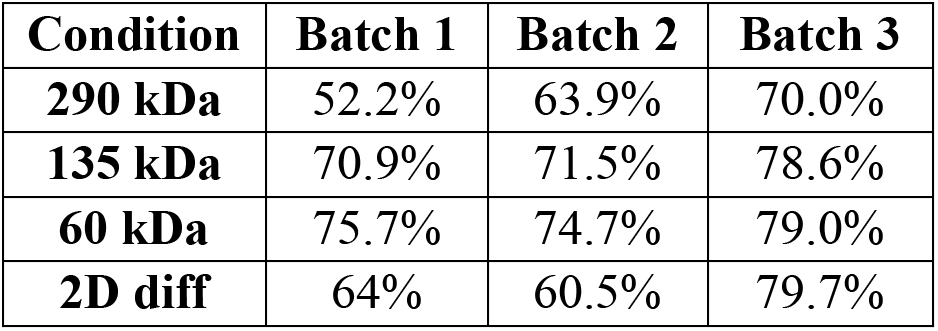
Cardiomyocyte purity across the batches used for µHT fabrication.

| **Condition** | **Batch 1** | **Batch 2** | **Batch 3** |
|---|---|---|---|
| **290 kDa** | 52.2% | 63.9% | 70.0% |
| **135 kDa** | 70.9% | 71.5% | 78.6% |
| **60 kDa** | 75.7% | 74.7% | 79.0% |
| **2D diff** | 64% | 60.5% | 79.7% |

Representative calcium transient and force traces illustrate the superior functional output of µHTs derived from hiPSC-CM derived in softer gels (**Figs. 6B, 6C**). Quantitative analysis of peak values and kinetics of calcium and force confirmed the significance of these functional differences.

Total calcium release, measured as area under the curve (AUC) of GCaMP fluorescence transients (**Supplementary videos 4-6**), was significantly higher in 135 kDa and 60 kDa EiC-derived µHTs compared to 290 kDa (**Fig. 6D**) and 2D monolayer differentiation conditions (**Fig. 7E**). Interestingly, µHTs fabricated from softer gel-derived cardiomyocytes generated significantly greater peak stress compared to µHT derived from hiPSC-CM from 290 kDa-gels (**Fig. 6E**). To determine whether these differences reflected intrinsic contractility, we measured peak active stress by normalizing peak active force to corresponding μHT cross-sectional area. Peak stress was significantly higher in 60 kDa EiC-derived µHTs compared to both 290 kDa and 135 kDa gels (Kruskal-Wallis test, p < 0.0001; Dunn’s post-hoc: 290 vs. 135 kDa, ns; 290 vs. 60 kDa, ****; 135 vs. 60 kDa, ****; **Fig. 6E**).

**Figure 7.**
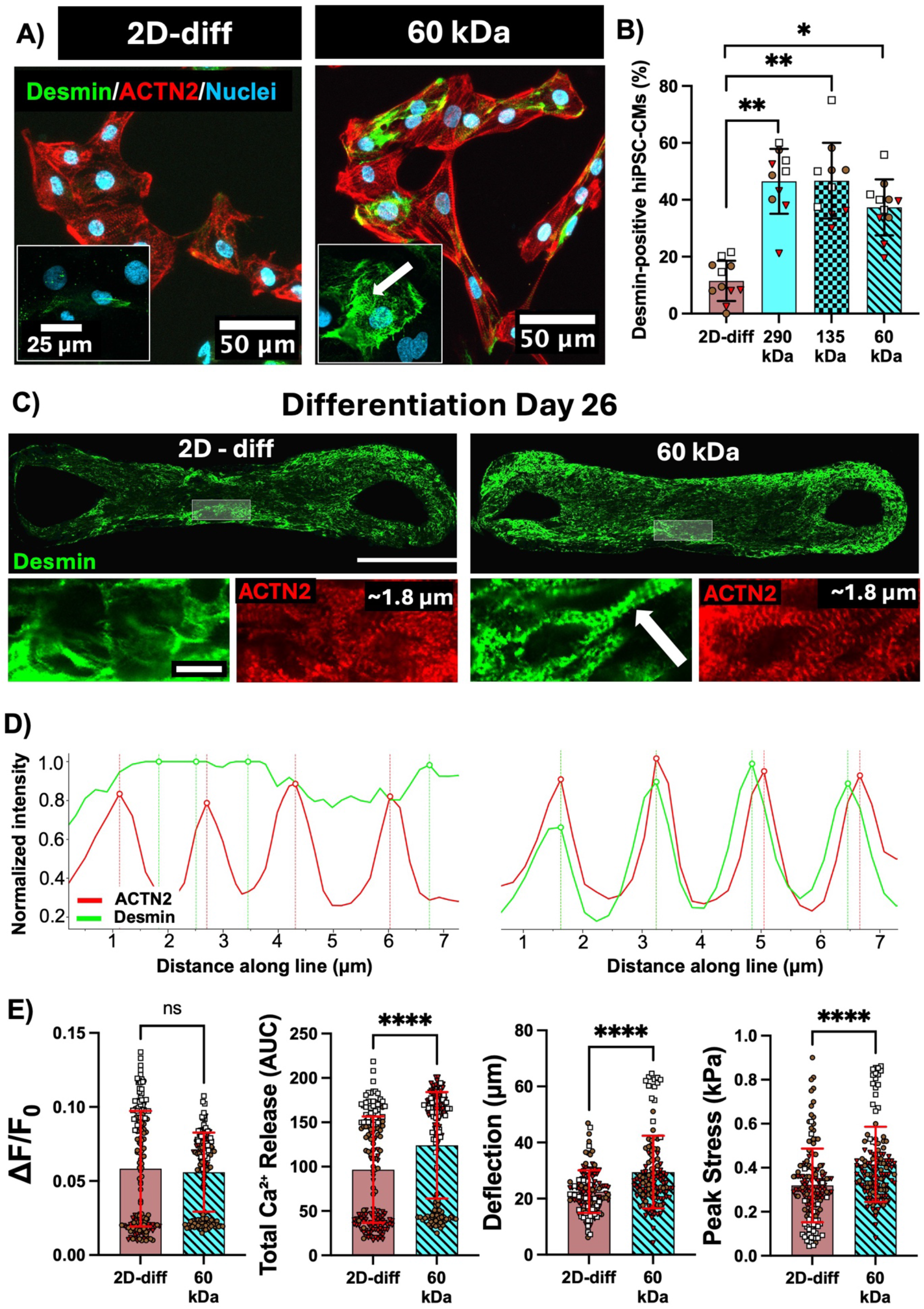
Desmin expression in 3D-differentiated hiPSC-CMs at day 15 and post-µHT fabrication. Softer gel-derived hiPSC-CMs exhibit improved desmin striation within µHTs. A) Representative immunofluorescence images of dissociated hiPSC-CMs at day 15 stained for Desmin (green), ACTN2 (red), and nuclei (cyan) across 2D monolayer differentiation and 60 kDa at 25 µM cRGD. High-magnification immunofluorescence images of Desmin (green) and nuclei (cyan) in dissociated hiPSC-CMs from 2D monolayer and 60 kDa conditions. Arrows indicate regions of striated Desmin organization in 60 kDa-derived cardiomyocytes, absent in 2D-derived cells. **B)** Desmin-positive hiPSC-CMs as a percentage of total cells at day 15; each dot represents one imaging region, colored by batch. N = 3 independent differentiation batches per condition (n = 11, 2D-diff; n = 10, 290 kDa; n = 10, 135 kDa; n = 11, 60 kDa). Nested one-way ANOVA with Tukey’s multiple comparisons test was used for B. 2D-diff vs. 290 kDa, **p = 0.0035; 2D-diff vs. 135 kDa, **p = 0.0040; 2D-diff vs. 60 kDa, *p = 0.0206; all other comparisons ns. **C**) Representative immunofluorescence images of Desmin (green) and ACTN2 (red) within intact µHTs fabricated from 2D monolayer- and 60 kDa-derived hiPSC-CMs at differentiation day 26. Insets show higher-magnification views; arrows highlight striated Desmin patterning, more prominent in the 60 kDa condition. Scale bars = 500 µm (whole µHT) and 10 µm (insets). **D)** Representative normalized fluorescence intensity line-scan profiles of ACTN2 (red) and Desmin (green) across hiPSC-CMs within µHTs, comparing 2D-diff and 60 kDa conditions. In the 60 kDa condition, Desmin intensity peaks align with ACTN2 peaks, indicating a striated, sarcomere-registered pattern; in 2D-diff, Desmin is expressed but does not show a periodic pattern coincident with ACTN2 striations. **E)** µHT functional measurements: calcium transient amplitude (ΔF/F₀), total Ca²⁺ release (AUC), deflection (µm), and Peak Stress (kPa) for 2D-diff and 60 kDa conditions. Bars represent mean ± SD (each dot represents one µHTs, pooled from N = 3 independent batches); ns = not significant, ****p < 0.0001.

To provide a comprehensive overview of functional differences, a panel of metrics including ΔF/F₀, UPD, AUC, peak active force, deflection, and work per cycle were normalized and visualized as a heatmap (**Fig. 6F**). This visualization consistently shows hiPSC-CM derived from softer gels outperformed hiPSC-CM derived from the stiffer 290 kDa gels, despite the functional analysis occurring nearly 2 weeks after retrieving the hiPSC-CM from gel-based EiC and culturing in a very similar µHT-based micro-environment. Taken together, these findings demonstrate that optimizing alginate matrices for hiPSC expansion and differentiation translates to meaningful improvements in the functional performance of downstream engineered cardiac tissues.

### hiPSC-CM derived from EiCs exhibit more “in vivo-like” desmin intermediate filament expression compared to 2D monolayer-derived hiPSC-CM

Prior transcriptomic analysis of hiPSC cardiac differentiation has demonstrated that 3D culture conditions accelerate cardiomyocyte maturation compared to conventional 2D monolayer differentiation, with Desmin (DES) specifically enriched in 3D aggregates by differentiation day 9, coinciding with the emergence of cardiac conduction system gene expression [11]. Importantly, Desmin’s role extends beyond serving as a passive structural marker. Desmin deficiency has been shown to actively impair cardiac maturation by diminishing the expression and proper localization of cardiac-specific proteins, disrupting calcium homeostasis, and delaying myofibril formation [42]. These observations suggest that robust Desmin expression and striated organization is not merely a readout of maturation but may itself be a prerequisite for functional cardiomyocyte development. Motivated by these findings, we examined Desmin expression and striated organization in hiPSC-CMs derived from our EiC system across alginate molecular weights, using 2D monolayer derived hiPSC-CM as a benchmark.

Consistent with our previous study [25] and work of other teams [7], we found that hiPSC-CM derived from standard 2D monolayers exhibited poor Desmin expression (**Fig. 7A**). In contrast, close to half of the hiPSC-CM derived from EiC exhibited prevalent Desmin expression (**Fig. 7B**).

Interestingly, however, no significant differences in Desmin positivity were detected among hiPSC-CM derived from gels with different mechanical properties. These findings indicate that 3D encapsulation in alginate promotes greater Desmin expression regardless of gel molecular weight, suggesting that the 3D microenvironment itself, rather than gel stiffness specifically, is the primary driver of improved structural maturation compared to conventional 2D differentiation. Higher magnification imaging further revealed qualitative differences in Desmin organization between conditions (**Fig. 7A**). While Desmin-positive cells were present in both 2D monolayer and 60 kDa EiC-derived cardiomyocytes, cells retrieved from 60 kDa EiCs displayed nascent striated Desmin patterning, indicated by arrows, consistent with the early organization of the intermediate filament network around developing sarcomeres. In contrast, 2D monolayer-derived cardiomyocytes showed a more diffuse, unorganized Desmin distribution, suggesting a lower degree of cytoskeletal maturation despite comparable culture duration.

In our prior studies, post-differentiation maturation of hiPSC-CM within engineered tissues proved to be a necessary step to observe appropriate expression of structural proteins, including Desmin [25, 26]. We thus sought to determine whether post-differentiation maturation in µHT could “cancel out” the apparently accelerated structural maturation of EiC-derived hiPSC-CM. For this analysis, we used µHT derived from “elite” 2D hiPSC-CM monolayers, which had high purity and produced µHT with superior contractility levels (**Fig. S6**). Whole-mount immunofluorescence staining for Desmin and ACTN2 within intact µHTs revealed progressively greater sarcomeric organization and more pronounced Desmin striation when the source hiPSC-CM were derived from softer gels, with 60 kDa EiC-derived µHTs showing the most organized striated pattern at this timepoint (differentiation day 26; **Fig. 7C and Fig. S6B**). Line-scan analysis of fluorescence intensity confirmed this qualitative impression: in the 60 kDa condition, Desmin intensity peaks aligned with ACTN2 peaks, indicating a striated, sarcomere-registered pattern, whereas in 2D-diff-derived µHTs Desmin was expressed but did not show a periodic pattern coincident with ACTN2 striations (**Fig. 7D**). Strikingly, despite our use of “elite” 2D monolayer differentiations as a benchmark, hiPSC-CM within µHT derived from 60 kDa EiC showed superior structural organization, including Z-disc localized Desmin, as compared to hiPSC-CM within µHT derived from the monolayers. Improved structural organization within the µHT formed from 60 kDa-derived cardiomyocytes is consistent with the functional improvements observed when µHT were formed from these gel-derived, rather than 2D monolayer-derived cardiomyocytes (**Fig. 7E**). Although background-corrected calcium transient amplitude (ΔF/F₀) was similar between conditions (ns), total calcium release (AUC) and deflection were both significantly higher in 60 kDa-derived µHTs, concurrent with significantly higher peak stress (**Fig. 7E**). Collectively, these findings support the conclusion that softer alginate matrices not only enhance cardiomyocyte yield but also promote a more structurally mature phenotype that translates into superior µHT functional performance.

Altogether, these findings demonstrate that 3D encapsulation in cRGD-modified alginate hydrogels not only facilitate differentiation but also promote cardiomyocytes’ structural maturation beyond what is achievable with conventional 2D monolayer differentiation. Reducing the stiffness of gels used to derive the hiPSC-CM enhances this effect, linking the biophysical properties of the encapsulation matrix during hiPSC expansion and differentiation to the structural and functional quality of the resulting hiPSC-CM, even with post-differentiation maturation in engineered cardiac tissue.

## Discussion

In this study, we developed a cRGD-functionalized alginate platform with independently tunable biochemical and mechanical properties for hiPSC encapsulation and *in situ* cardiac differentiation. Our findings reveal that both cRGD ligand density and gel stiffness are critical, independently acting determinants of hiPSC fate within the encapsulation matrix, and that their joint optimization yields structurally mature cardiomyocytes with functional properties comparable to those of tissue derived from the highest quality 2D monolayer-based differentiations.

cRGD functionalization proved essential for hiPSC survival and pluripotency maintenance within alginate hydrogels. In its absence, cells rapidly lost viability and Nanog expression within days of encapsulation, underscoring the fundamental incompatibility of bioinert alginate with pluripotent stem cell survival in the absence of integrin-mediated adhesion signaling [30]. Critically, however, 25 µM cRGD was sufficient to achieve maximal hiPSC viability, pluripotency, and metabolic support, with no additional benefit conferred by 75 µM, potentially indicating integrin occupancy saturation at this relatively low ligand density. Interestingly, this effective dose is 60-fold lower than RGD concentrations previously reported to support human pluripotent stem cell maintenance in a similar alginate system [30]. We attribute this finding to two factors: 1) our use of cyclic RGD peptides, which have been observed to be ∼10-fold-more potent than linear RGD in promoting stem cell survival and fate decisions [43], and 2) the SPAAC-based bioconjugation chemistry we employed here, in which cRGD is tethered to the alginate backbone via a BCN heterobifunctional spacer that substantially improves peptide accessibility and integrin engagement relative to conventional carbodiimide coupling strategies [23].

Beyond biochemical functionalization, gel mechanical properties exerted a profound influence on hiPSC behavior prior to and during cardiac differentiation. Softer 60 kDa gels supported greater colony spreading, rounder and more defined colony morphologies, and higher metabolic activity compared to stiffer 290 kDa gels, despite equivalent cRGD density. The qualitative observation of lumen-like structures within colonies in softer gel conditions is particularly intriguing, suggesting that reduced matrix resistance may facilitate early self-organizational behavior of hiPSCs analogous to that observed in compliant natural matrices [30]. These morphological differences likely reflect differences in the ability of hiPSCs to generate and transmit cytoskeletal tension against matrices of differing stiffness, and potentially to produce long-range mechanical deformation to facilitate colony expansion. In stiffer gels, constrained colony expansion and elevated compressive resistance may limit the cytoskeletal reorganization required for optimal pluripotent expansion [44, 45]. The temporal evolution of colony area (**Fig. 3A, B**) showed that softer 60 kDa gels supported greater colony expansion over the course of hiPSC culture compared to stiffer 290 kDa gels, consistent with the elevated metabolic activity and higher final cell counts observed in these same softer gels (**Fig. 3F**). The higher metabolic activity observed in 60 kDa gels at day 0 normalized to day −2, together with the directly measured higher cell number (**Fig. 3F**), confirms that softer gels support greater hiPSC expansion rather than simply reflecting higher per-cell metabolic activity. Together, these converging measures of greater colony expansion, higher metabolic activity, and higher cell number in softer gels provide a consistent explanation for the higher cardiomyocyte yield subsequently observed in 60 kDa-derived EiCs (**Fig. 4H**), linking enhanced hiPSC expansion capacity directly to downstream cardiomyocyte production. These observations are consistent with an emerging body of work demonstrating that matrix mechanical properties, including not only stiffness but also viscoelastic character, can independently govern stem cell behavior and differentiation outcomes. For example, viscoelastic, stress-relaxing hydrogels have been shown to support cardiomyocyte spreading, phenotypic maintenance, and functional maturation in ways that purely elastic matrices of equivalent stiffness cannot [46]. We note that in our prior study, we observed that lowering alginate M_w_ diminished stiffness *and* enhanced stress-relaxation (e.g. lower M_w_ gels exhibited shorter stress-relaxation times [24]) – it is possible that this stress-relaxation is the predominant reason why we observe superior hiPSC expansion in the softer, 60 kDa alginate-derived gels.

The improvements in hiPSC expansion and metabolic activity observed in softer gels translated directly into enhanced cardiomyocyte production. EiCs derived from 60 kDa gels showed a trend toward higher calcium transient amplitudes and significantly greater cardiomyocyte purity compared to EiCs derived from 290 kDa gels.

EiCs derived from 60 kDa gels exhibited slightly higher calcium transient amplitudes and greater cardiomyocyte purity compared to EiC derived from 290 kDa gels, a finding that initially appeared to suggest differences in per-cell calcium handling. However, equalization of ΔF/F₀ and upstroke duration upon matching for cardiomyocyte content across conditions reveals that these differences are attributable entirely to differences in CM yield rather than intrinsic electrophysiological properties. This demonstrates that softer matrices enhance cardiomyocyte production efficiency without altering the fundamental calcium handling capacity of individual cells.

The significantly higher cTnT⁺ purity and yield in 60 kDa gels points to matrix molecular weight as a determinant of differentiation efficiency, likely reflecting the combined effects of improved hiPSC expansion, and more favorable matrix remodeling during cardiogenesis. The precise mechanisms by which gel mechanics influence cardiomyocyte output remain to be established; however, integrin-mediated mechanotransduction represents a plausible upstream contributor, given prior evidence that matrix physical properties modulate the signaling pathways governing early mesodermal commitment [47–49]. Elucidating these mechanisms will require future studies with direct perturbation of mechanosensing pathways at defined stages of differentiation. Notably, comparison with established 2D monolayer and 3D suspension differentiation platforms (**Table S2**) indicates that the pre-differentiation hiPSC expansion window itself is a yield-determining parameter: published protocols span Day −2 to Day −5 of expansion prior to initiating differentiation, and in at least one system a shorter expansion window (Day −2) outperformed a longer one (Day −4), indicating that more expansion time does not uniformly improve yield. Nonetheless, for a given protocol, including ours, it is likely that increasing the timeframe of hiPSC expansion could further increase the ratio of cardiomyocytes yielded per initial hiPSC seeded.

In the present study, we used a single, non-optimized Day −3 expansion window across all conditions; systematic optimization of this parameter for the EiC platform represents a straightforward, unexplored opportunity to further increase cardiomyocyte yield.

A notable finding of this study is the demonstration that 3D encapsulation in cRGD-modified alginate, regardless of molecular weight, dramatically increases the proportion of Desmin-positive cardiomyocytes compared to 2D monolayer differentiation. The low proportion of Desmin-positive cells observed in 2D monolayer differentiation in the present study (11.5%) is consistent with prior reports of limited desmin expression in hiPSC-CMs cultured under standard 2D conditions (∼10%) [50], validating our 2D baseline and underscoring the inherent limitations of monolayer culture in supporting intermediate filament network maturation. This is further consistent with prior transcriptomic evidence that 3D culture conditions enrich for Desmin expression during cardiac differentiation [11]. Notably, this enhancement appears to be a feature of the 3D microenvironment itself rather than a response graded by the stiffness of the matrix used to encapsulate hiPSC, at least over the range of stiffness we used in this study. This implicates encapsulation-driven mechanical and adhesive cues as key regulators of intermediate filaments. This may prove particularly valuable for modeling Desmin-Related Cardiomyopathy, where the absence of proper Desmin localization in conventionally differentiated hiPSC-CMs has been a fundamental barrier to developing human cell-based disease models and testing targeted therapies. We note the higher baseline from 2D differentiation we observed here, compared to our prior study [25], may reflect that in that prior work, we dissociated 2D monolayer differentiations to replate hiPSC-CM for purification in lactate-enriched, glucose depleted media, and this extended culture in a sparse 2D monolayer may have diminished an already low baseline-level of Desmin expression.

Interestingly, gel stiffness did exert an influence on the quality of Desmin organization within intact µHTs, with softer 60 kDa-derived µHTs exhibiting the most pronounced striated Desmin patterning, progressively more organized than 135 kDa, 290 kDa, and 2D-derived µHTs. This gradient of structural organization mirrors the stiffness-dependent improvements in contractile force and calcium handling observed across the same conditions, suggesting that the mechanical environment during differentiation establishes a lasting epigenetic or structural memory that influences cardiomyocyte organization within downstream tissue constructs. The lower MLC2a expression, higher MYH7 expression, and greater cell area and lower cell circularity of 60 kDa EiC-derived cardiomyocytes compared to 2D controls further support the conclusion that softer matrices accelerate structural maturation beyond what is achievable in conventional monolayer differentiation, consistent with prior transcriptomic comparisons of 3D versus 2D cardiac differentiation [11]. Potential impacts of epigenetic imprinting during early stages of hiPSC differentiation to cardiomyocytes have been noted by prior teams. For example, Dark *et al.* describe how optimizing differentiation conditions at the mesoderm stage produces superior maturation in left-ventricular-like hiPSC-CM [51]. Similarly, Krup *et al.* demonstrated that expression of the early mesodermal marker MESP1 is crucial to eventually generate mature cardiomyocytes *in vivo* [52].

The functional benefits of EiC-based differentiation were clearly demonstrated in the µHT platform, where cardiomyocytes from softer gel conditions produced significantly higher peak active force and total calcium release compared to stiffer conditions and 2D-derived controls. The improvement in contractile output was stiffness-dependent: hiPSC-CM derived from 60 kDa EiC formed µHTs that outperformed those made from 135 kDa and 290 kDa EiC. These findings represent a remarkable translational link between the biophysical properties of the encapsulation matrix and the functional quality of downstream engineered cardiac tissues, even when these hiPSC-CM are encapsulated into a common pro-maturation engineered tissue environment and cultured there for an extended timeframe. The relatively similar cardiomyocyte purity across the differentiation batches we used for µHT fabrication (**Table 2**) suggests that these functional differences are intrinsic to the cells themselves rather than a consequence of differential CM content. The observation that hiPSC-CM derived from 60 kDa and 135 kDa gels produced µHT that performed equally as well, if not better than hiPSC-CM even from µHT made from hiPSC-CM derived from optimized 2D monolayer-differentiations is particularly significant, as it suggests that the functional benefits of 3D gel-based differentiation are not merely equivalent to, but in some cases may exceed what is achievable through conventional approaches, even under optimized conditions. EiC-derived hiPSC-CM also produced µHTs with significantly higher peak contractile stress than those derived from elite 2D monolayer differentiations, with slightly lower µHT-to-µHT variability and evidence of improved structural maturation.

Our platform sits within a broader effort to differentiate hiPSCs into cardiomyocytes within 3D matrices. Several studies over the past decade have pursued *in situ* cardiac differentiation, in which hiPSCs differentiate while embedded within a 3D matrix rather than being encapsulated after differentiation. Early work established feasibility using gelatin-based materials. Kerscher et al. [53] and Kupfer et al. [54] demonstrated this using GelMA and ECM-protein bioinks, respectively; the latter showed that allowing hiPSCs to proliferate to tissue-like densities before differentiation could overcome the poor proliferation and migration of cardiomyocytes and yield contiguous muscle. These studies suggest that coupling differentiation to 3D tissue formation is important for *in situ* differentiation to succeed. However, both of these prior foundational works relied on animal-derived gelatin or native ECM proteins, which inevitably leads to batch variability, along with challenges, including on a regulatory level, stemming from the materials being incompletely defined.

Subsequent studies using Matrigel-based hydrogel systems further demonstrated the fidelity of this approach. Pramanick et al. [55] demonstrated *in situ* differentiation within shape-morphing Matrigel constructs, reporting that cardiomyocytes and fibroblasts co-emerge from a common progenitor pool and that support-bath mechanics influence iPSC fate. Banerjee et al. [56] observed a similar effect in a fish-gelatin GelMA hydrogel (FiBGel), where *in situ* differentiation of encapsulated hiPSCs produced organized, marker-positive, contractile tissue that remained functional for long timeframes.

More recent work has pursued more defined formulations, overcoming batch variability and xenogeneic components of animal derived ECM proteins or ECM protein mixtures like Matrigel. Ong et al. [18] addressed the materials problem directly, developing a clinically defined, xeno-free fibrin-laminin system that supports trilineage differentiation, including cardiac, although fibrin remains a human blood-derived product. A related approach used PEG-fibrinogen, a hybrid synthetic-natural biomaterial, to move past the variability of Matrigel. Encapsulating hiPSCs in PEG-fibrinogen microspheres, Hashemi et al. [12] reported that *in situ* differentiation outperformed scaffold-free aggregates, the current standard for scaling hiPSC-CM production, yielding higher cardiomyocyte content and yield, improved functionality, and greater batch-to-batch consistency. These results suggest that even a relatively simple engineered niche can outperform scaffold-free aggregates on both quality and reproducibility. However, the bovine fibrinogen component, like Ong’s human fibrin, only partially resolves the definability problem.

A parallel line of work has pursued fully defined, peptide-functionalized hydrogels as xeno-free alternatives to Matrigel, addressing the definability limitation noted above. Several groups have used RGD-functionalized PEG substrates with hiPSCs: Mulero-Russe et al. [57] supported definitive endoderm differentiation, and Amitrano et al. [58] identified cRGD-PEG formulations that improved the efficiency and reproducibility of iPSC-cardiomyocyte differentiation over Matrigel. In both cases, however, cells were cultured as a 2D monolayer on the gel surface rather than encapsulated. Other defined systems have achieved 3D hiPSC encapsulation but in different contexts: for example, Li et al. [59] used a self-assembling peptide hydrogel to encapsulate hiPSCs and direct *in situ* hepatocyte differentiation.

Across this body of work is a key recurring theme our current work builds on: the matrix instructs cell fate through both its mechanics and its adhesive ligands, and a persistent limitation has been achieving these outcomes in a fully defined, animal-free material. The cRGD-alginate system addresses the latter directly, replacing the animal-derived protein matrices used in most prior in situ cardiac differentiation work. It also addresses the former: alginate stiffness and stress relaxation can be tuned through molecular-weight blending independently of cRGD density, decoupling matrix mechanics from biochemical adhesion. These two cues remain coupled in protein-based systems, where stiffness, composition, and ligand content tend to co-vary. The EiC platform therefore provides a fully defined, xeno-free material in which adhesive ligand and matrix mechanics can be set independently. To our knowledge, no prior study has combined a defined, peptide-functionalized matrix with hiPSC encapsulation and *in situ* cardiac differentiation.

Collectively, our findings establish an encapsulation platform that provides several features that make it highly desirable for iPSC-CM biomanufacturing, including improved batch-to-batch reproducibility, enhanced cardiomyocyte yield, and accelerated structural maturation, while preserving the ability to execute complex downstream applications such as tissue engineering. Because these outcomes are achieved in a chemically defined, animal-free material, the platform is well positioned for the xeno-free requirements of clinical translation and regulatory approval [18], and its tunability through simple formulation adjustments, without changes to total polymer concentration or crosslinking chemistry, offers a flexible and scalable design framework.

Several directions will be important for future study. First, the mechanistic basis for stiffness-dependent changes in hiPSC-CM differentiation and maturation within EiCs remains to be established. Potential roles for mechanosensitive transcription factors such as YAP/TAZ will be important to pursue. It would also be informative to directly compare hiPSC-CM differentiated as a 2D monolayer atop cRGD-functionalized alginate substrates to those differentiated *in situ* within 3D cRGD-alginate encapsulation, which would help isolate the contribution of 3D geometry itself from that of matrix composition and stiffness. A limitation of the present study is that our assessment of cardiomyocyte maturation relied on protein- and structural-level markers; transcriptomic, electrophysiological, and mitochondrial metabolic characterization were not performed here. Future work incorporating transcriptomic profiling, such as bulk or single-cell RNA sequencing, would provide additional information on how transcriptional shifts relate to protein-level structural maturation. Likewise, analysis of cardiomyocytes’ metabolic state and patch-clamp analysis of ion channel activity and resting membrane potential will be important future steps to assess the function of EiC-derived iPS-CM. Functionally, µHTs made from EiC derived hiPSC-CM had contractile performance that *exceeded* the performance of µHT made from optimized monolayer-derived hiPSC-CM. This structural and functional maturation may be enhanced further by dynamic mechanical stimulation and/or prolonged culture on cardiomyocyte maturation within these gels, both of which warrant further study. Finally, integration of this platform with metabolic purification strategies and large-scale suspension bioreactors will be a critical step toward clinical-scale hiPSC-CM production.

## Conclusion

We developed a xeno-free, cRGD-functionalized alginate platform that encapsulates hiPSCs and supports their expansion and *in situ* cardiac differentiation within a fully defined 3D matrix. By decoupling adhesive ligand density from matrix mechanics, two cues that remain coupled in protein-based systems, we showed that each independently shapes hiPSC fate: cRGD functionalization is required for survival and pluripotency, while softer, lower-molecular-weight gels enhance expansion, cardiomyocyte yield, and structural maturation. Cardiomyocytes generated within this 3D environment exhibited signs of structural maturation, including markedly higher Desmin content, than those from 2D monolayer differentiation. When assembled into micro heart tissues, hydrogel-derived cardiomyocytes yielded tissues with significantly higher peak contractile stress than those derived from elite 2D monolayer differentiation. These outcomes were achieved in a chemically defined, animal-free material amenable to simple formulation-based tuning, addressing a central barrier to the clinical translation of hiPSC-CMs. Together, this work establishes independently tunable matrix adhesion and mechanics as a practical design framework for producing structurally more mature hiPSC-derived cardiomyocytes in a scalable, fully defined manner.

### Experimental Section

#### Human iPSC culture

Human induced pluripotent stem cells (hiPSCs) (Wild-type C, WTC; Coriell Repository # GM25256) harboring a single-copy of CAG-driven GcaMP6f [60]) were a kind gift from Dr. Bruce Conklin, and were obtained under MTA from the Gladstone Institutes of Cardiovascular Disease. hiPSC were cultured in StemFlex media (Life Technologies; A3349401) on Geltrex-coated (Life Technology; A1413302) tissue culture plates at a concentration of 18.75µg/cm^2^. During passaging, hiPSCs were rinsed with Dulbecco’s Phosphate Buffered Saline (dPBS) and dissociated as single cells using Gentle Dissociation Reagent (dPBS with 1.8 g/L NaCl and 0.5 mM EDTA [61]) and replated in media supplemented with 10 µM Y-27632 (ROCK inhibitor; Biogems). The next day, media was replaced with Stemflex without Y-27632. Media was changed daily and cells were passaged every 3 days at a density of 30,000 cells/cm^2^.

### cRGD conjugation to alginate

Alginate was functionalized with cyclo[Arg-Gly-Asp-D-Phe-Lys(azide)] (cRGD) adhesion peptides via strain-promoted azide-alkyne cycloaddition (SPAAC) bioconjugation, linking azide-terminated cRGD to BCN-modified alginate through a triazole linkage (**Fig.1A**). BCN-modified alginate was prepared as described previously [23, 62]. Briefly, high molecular weight alginate (Manugel, Dupont) was dissolved at 1% wt/v in 0.1 M (2-(N-Morpholino)ethanesulfonic acid, 4-Morpholineethanesulfonic acid) (MES) buffer (pH 6.0). *N*-[(1*R*,8*S*,9*s*)-Bicyclo[6.1.0]non-4-yn-9-ylmethyloxycarbonyl]-1,8-diamino-3,6-dioxaoctane (BCN-amine; Sigma-Aldrich), *N*-(3-Dimethylaminopropyl)-*N*′-ethylcarbodiimide (EDC; ThermoFisher), and *N*-Hydroxysuccinimide (NHS; ThermoFisher) were mixed in MES buffer and added dropwise to the stirring alginate solution at a 20-fold molar excess of BCN-amine, 36-fold molar excess of EDC, and 7-fold molar excess of NHS relative to BCN-amine, and allowed to react for 24 hours. The solution was precipitated twice in a 10-fold volumetric excess of methanol, dried under vacuum, and resuspended in deionized water for lyophilization. cRGD-azide peptides (Biosynth) were then click-conjugated to BCN-alginate via SPAAC reaction (24 hr in dPBS). Following conjugation, peptide-modified alginates were dialyzed extensively, sterile filtered (0.2 µm), and lyophilized.

### Controlling alginate molecular weight

Alginate molecular weight (M_w_) was reduced by autoclaving high molecular weight alginate (290 kDa, Manugel) to achieve target molecular weights, as described previously [24]. Briefly, alginate polymers were subjected to repeated autoclaving cycles (10 minutes at 121°C) to reduce M_w_ from 290kDa (baseline) to ∼135 kDa (1 cycle) or ∼60 kDa (2 cycles). These molecular weights were previously verified by gel permeation chromatography [24]. Following M_w_ reduction, alginates were dialyzed extensively, sterile filtered and lyophilized. This autoclaving treatment does not alter alginate chemical composition or Ca²⁺-crosslinking capacity: we previously confirmed via ¹H-NMR and FTIR that the G/M ratio and chemical structure of alginate are unchanged across autoclave cycles, and that gelation kinetics and equilibrium swelling of Ca²⁺-crosslinked hydrogels are unaffected by the number of autoclave cycles, while hydrogel mechanical behavior change systematically with molecular weight [24].

### Mechanical properties characterization

Alginate polymers were dissolved into dPBS. Polymers were combined with calcium carbonate and GDL to final concentrations of 1% wt/v alginate with 22.5 mM CaCO₃ and 45 mM GDL and dispensed into disk shaped molds with a diameter of 10 mm diameter and height of 3 mm and allowed to crosslink for 45 minutes [63]. Next, the gels were equilibrated in Dulbecco’s Modified Eagle Medium (DMEM) for 24 hours at 37° C. Compressive elastic modulus was measured using an ElectroForce 3200 mechanical testing system (TA Instruments) with a 10 N load cell, compressing samples from 0–10% strain at 1 mm/min.

### Human iPSC encapsulation in alginate hydrogels

All hydrogels were prepared at a final alginate concentration of 1% wt/v. cRGD-modified 290 kDa alginate (stock concentration: 150 µM [23]) was blended with unmodified alginate of the desired molecular weight (290, 135, or 60 kDa) to achieve the target cRGD concentration, while the bulk mechanical properties of the hydrogel were governed by the molecular weight of the unmodified alginate component (**Table 1)**. hiPSCs were dissociated into single cells using Gentle Dissociation Reagent, pelleted by centrifugation, and resuspended in StemFlex medium supplemented with 10 µM Y-27632. CaCO₃ was added to the cell suspension and mixed thoroughly, followed by addition of the alginate solution and gentle mixing. Filter-sterilized GDL solution was then added to initiate crosslinking, achieving final concentrations of 4 × 10⁶ cells/mL, 1% wt/v alginate, 22.5 mM CaCO₃, and 45 mM GDL. The pre-hydrogel mixture was dispensed into PDMS molds (5 mm diameter, 1 mm thickness) placed in a 6-well plate and allowed to crosslink for 30 to 45 minutes at 37°C. Encapsulated hiPSCs were transferred to ultra-low adhesion 6 well plates and cultured in StemFlex medium supplemented with 10 µM Y-27632. The next day (day -2), medium was changed to Stemflex without Y-27632. Media was again changed on day -1. The following day (day 0) marked the time either for initiating cardiomyocyte differentiation or assessing hiPSC viability and pluripotency.

### Human iPSC-laden gel geometry characterization

To characterize hiPSC colony morphology within alginate hydrogels, brightfield images were acquired at designated timepoints using a Nikon Ts2R-FL inverted microscope. For colony area quantification, cell-laden hydrogels were fixed and stained with Alexa Fluor 633 Phalloidin (ThermoFisher) to visualize F-actin. Briefly, hiPSC-laden alginate hydrogels were washed twice with 100 mM HEPES supplemented with 2 mM CaCl₂ (HEPES/CaCl₂) and fixed in HEPES/CaCl₂ containing 4% paraformaldehyde for 20 minutes on a shaker plate. Gels were washed twice with HEPES/CaCl₂ and permeabilized with 0.1% Triton-X-100 for 30 minutes, repeated once for a total permeabilization time of 1 hour. Following permeabilization, gels were blocked in 2.5% BSA and 2.5% normal goat serum for 4 hours. Primary antibodies were added and incubated for 72 hours at 4°C on a shaker plate. Primary antibodies used include rabbit anti-Nanog (Cell Signaling Technology). Gels were washed three times with 0.1% Triton-X-100 and incubated with secondary antibodies (Alexa Fluor 488 or 594, ThermoFisher), Phalloidin (Alexa Fluor 633, ThermoFisher), and Hoechst 33342 (ThermoFisher) for overnight. Imaging was performed by Fluoview FB1200 confocal microscope (Olympus, Tokyo, Japan). Individual colony boundaries were identified using automated thresholding in ImageJ, and colony area was quantified from the projected images.

### Metabolic activity assessment and hiPSC viability

Metabolic activity of hiPSC-laden alginate hydrogels was assessed using resazurin reduction (Alamar Blue cell viability reagent; ThermoFisher). At designated timepoints, cell-laden hydrogels were transferred to a fresh ultra-low adhesion plate and incubated with Alamar Blue solution (10% v/v in phenol red-free DMEM (Gibco; 31053-028) supplemented with sodium pyruvate) for 4 hours at 37°C. Fluorescence of the reduced product was measured on a plate reader (ex: 560 nm, em: 590 nm). Metabolic activity at day 0 was normalized to the day −2 baseline measurement for each tissue to account for differences in initial cell seeding. Viability of hiPSC-laden alginate hydrogels was evaluated using a LIVE/DEAD Viability/Cytotoxicity Kit (Invitrogen). Tissues were washed with dPBS for 5 minutes, incubated with the live/dead staining solution for 20 minutes at room temperature on shaker, and immediately imaged using a fluorescence microscope (Nikon). Image analysis was performed using ImageJ. Cell viability following gel dissociation was assessed using a Zombie dye viability kit (BioLegend) according to the manufacturer’s instructions and analyzed by flow cytometry as described below.

### 2D monolayer-based hiPSC-CM differentiation

hiPSCs were seeded onto Geltrex-coated tissue culture plates at approximately 26,000 to 32,000 cells/cm² and cultured in StemFlex medium until reaching 80 to 90% confluency, marking Day 0. Cardiac differentiation was then initiated using the GiWi protocol [2, 31]. Briefly, on day 0, medium was switched from StemFlex to RPMI 1640 supplemented with B27 minus insulin (ThermoFisher) and 150 µg/mL ascorbic acid, and cells were treated with 6 µM CHIR99021 (Biogems) for 48 hours to activate Wnt signaling at 37°C with 5% CO₂. On day 2, CHIR99021-containing medium was replaced with fresh RPMI 1640 supplemented with B27 minus insulin, 150 µg/mL ascorbic acid, and 5 µM IWP2 (Biogems) to inhibit Wnt signaling for an additional 48 hours. On day 4, fresh RPMI 1640 supplemented with B27 minus insulin and 150 µg/mL ascorbic acid was added. From day 6 onward, cells were maintained in RPMI 1640 supplemented with B27 plus insulin (ThermoFisher), with medium changes every 48 hours. Cardiomyocyte differentiation was assessed at day 15.

### 3D EiC (in-gel) hiPSC-CM differentiation

Cardiac differentiation of hiPSC-laden alginate hydrogels to form “Engineered iPSC Cardiobodies” (EiC) was initiated at Day 0 (defined above) using the same GiWi protocol, small-molecule concentrations, and exposure durations described for 2D monolayer differentiation above. Briefly, cells were treated with 6 µM CHIR99021 for 48 hours followed by 5 µM IWP2 for 48 hours, with all culture steps performed at 37°C on an orbital shaker (50 ± 2 rpm) with 5% CO₂ to keep hydrogels in suspension. Cardiomyocyte differentiation was assessed at day 15.

### Micro heart tissue (µHT) formation and culture

Following cardiac differentiation, hiPSC-CMs were retrieved from alginate hydrogels by gel dissociation as described in tissue dissociation section on day 15. Dissociated cells (including hiPSC-CMs) were resuspended at a total cell density of 2 × 10⁷ cells/mL in a mixture of 1.5 mg/mL collagen type I rat tail (Fisher Scientific; A1048301) and 15µg/mL Geltrex, and 4 µL of this mixture (80,000 cells total) was seeded into each 2-post µHT device. Devices were incubated at 37°C with 5% CO₂ for 20 minutes to allow collagen crosslinking, after which the culture well was filled with µHT formation media consisting of KnockOut DMEM supplemented with 20% Fetal Bovine Serum (FBS), 1% Glutamax, 1% non-essential amino acids, 1% penicillin-streptomycin, 150 µg/mL ascorbic acid, and 10 µM Y-27632. Tissue compaction was typically observed within 48 hours. After two days, media was changed to KnockOut DMEM supplemented with 1% KnockOut Serum Replacement, 1% Glutamax, 1% non-essential amino acids, 1% penicillin-streptomycin, and 150 µg/mL ascorbic acid, and refreshed every two days thereafter.

### Engineered iPSC Cardiobodies (EiCs) dissociation and 2D replating

To retrieve hiPSC-derived cardiomyocytes from alginate hydrogels, cell-laden gels were dissociated using a protocol adapted from Breckwoldt et al. [64]. Briefly, cell-laden gels were incubated with Collagenase II solution (200 units/mL in HBSS; Worthington, LS004176) supplemented with 10 µM Y-27632 and 30 µM N-benzyl-p-toluenesulfonamide (BTS) at 37°C on an orbital shaker (50 ± 2 rpm) for a minimum of 16 hours. Following incubation, tissues were gently triturated using a P1000 pipette to assess dissociation; if cell clumps remained, an additional 1 hour of incubation was performed. The collagenase reaction was quenched by addition of DNase wash buffer (1% BSA in DMEM/F-12 supplemented with DNase; MilliporeSigma, 260913-10MU) at 1–1.5× the volume of the collagenase solution. The cell suspension was transferred to centrifuge tubes and pelleted at 300 rcf for 5 minutes.

As collagenase digestion alone may not be sufficient to fully singularize hiPSC-CMs, an additional dissociation step was performed. Following centrifugation, the cell pellet was resuspended in TrypLE Select (1X; Gibco) and incubated at 37°C for 10 minutes to achieve single-cell dissociation. The reaction was quenched with an equal volume of EB20 medium (KnockOut DMEM supplemented with 10% FBS, 1% MEM/NEAA, and 1% GlutaMAX), and cells were pelleted again at 300 rcf for 5 minutes. The pellet was resuspended in RPMI/B27 plus insulin supplemented with 10% FBS and 10 µM Y-27632, triturated gently to ensure a homogeneous single-cell suspension, and filtered through a 70 µm cell strainer prior to cell counting.

For 2D replating, dissociated cells were seeded at approximately 4 × 10^4^ cells/cm² onto coverslips coated with 18.75 μg/cm^2^ Geltrexin RPMI/B27 plus insulin supplemented with 10 µM Y-27632 and 10% FBS. Cells were allowed to attach and spread for 48–72 hours prior to fixation and staining.

### Immunofluorescent visualization

For replated cells, dissociated cardiomyocytes on Geltrex-coated chamber slides (Millipore Sigma) were fixed in 4% paraformaldehyde in dPBS for 20 minutes at room temperature, permeabilized with 0.1% Triton-X-100 for 10 minutes three times, and blocked in 2.5% BSA and 2.5% normal goat serum for 2 hours. Cells were stained for and mouse anti-ACTN2 (Sigma-Aldrich) and MLC2a (SYSY) following the same secondary antibody protocol as above. To enhance myofilament visualization for Figure 5A inset images, an unsharp mask with weight 0.8 was applied in ImageJ.

### Automated image segmentation and cell morphology analysis

Confocal image stacks were processed using a custom Python pipeline. Each file was split into its constituent channels (DAPI, ACTN2, and the target protein, e.g., Desmin) and converted to single-plane images by maximum-intensity Z-projection. Stacks spanned approximately 10 µm for replated 2D monolayer cells and approximately 100–200 µm for 3D whole-mount stained samples.

Cell and nuclear segmentation were performed using Cellpose [65]. To segment individual cells, we leveraged a cyto2 model-based approach previously developed by our team [25], wherein the cortical component of the ACTN2 stain is used to delineate cell boundaries. Next, a Cellpose nucleus model was applied to the DAPI channel. Each identified nucleus was then assigned to the appropriate cell boundary (the cell boundary with which the given nucleus shared the greatest number of overlapping pixels). Only cells meeting predefined quality criteria (a minimum cell area of 96 µm^2^, an assigned nucleus, and a resolvable cytoplasmic region) were retained for analysis.

For each retained cell and nucleus, morphological parameters were quantified, including area, elongation (defined as the ratio of the major to minor axis of the best-fit ellipse), eccentricity, and circularity (defined as 4π × area/perimeter²). Target protein expression (e.g., Desmin) was quantified by measuring mean fluorescence intensity within the whole-cell, nuclear, and cytoplasmic compartments of each segmented cell. Cells were classified as protein-positive when their mean intensity met or exceeded a user-defined threshold (same threshold used for all images of the same protein antigen).

### Flow cytometry

Cardiomyocyte purity was assessed by flow cytometry using antibodies against cardiac troponin T (cTnT). Following gel dissociation as described above, single-cell suspensions were stained with Zombie Violet Fixable Viability Dye (BioLegend) to distinguish live from dead cells. Cells were then washed with dPBS and fixed and permeabilized overnight at 4°C using the eBioscience FOXP3/Transcription Factor Staining Buffer Set (Invitrogen) to enable intracellular antibody access. The following day, cells were incubated with primary antibodies for 1 hour at room temperature, washed, and incubated with appropriate fluorochrome-conjugated secondary antibodies for 45 minutes. Data were acquired on a NovoCyte^®^ 3000 flow cytometer and analyzed using FlowJo software.

### Optocardiography

Calcium dynamics in EiCs and µHTs were assessed by GCaMP6f fluorescence imaging. EiCs or µHTs expressing GCaMP6f were placed in a temperature-controlled imaging chamber at 37°C and electrically paced at 1 Hz using a field stimulator. Time-lapse fluorescence images were acquired at 100 frames per second using the Ts2R-Fl inverted fluorescence microscope equipped with a GFP filter set. Calcium transient parameters were extracted from regions of interest (ROIs) drawn over individual beating areas using custom analysis scripts in MATLAB [66]. Parameters quantified include calcium transient amplitude (ΔF/F₀), deflection (UPD, time from baseline to peak fluorescence), and decay time to from peak to 25% of peak amplitude (Decay 75%, τ_75_). Raw GCAMP fluorescence signals were background-corrected by subtracting the mean fluorescence intensity of a cell-free region of interest. Normalized fluorescence changes were calculated as ΔF/F0, where F0 represents the mean baseline fluorescence prior to calcium transient initiation. Total calcium release was quantified as the area under the curve (AUC) of the fluorescence transient. A minimum of 46 ROI recordings from at least three independent EiCs across three independent encapsulation and differentiation experiments were analyzed per condition.

### Contractile force quantification

Contractile force of µHTs was quantified from displacement of the flexible PDMS posts, as described in our prior work [25]. Bright-field videos were acquired at 100 frames per second. µHTs were electrically paced at 1 Hz with 20 msec bipolar pulses (<20 V; IonOptix Myopacer) by graphite electrodes (MyoPacer, Ion Optix, USA) during high-speed bright-field video acquisition. Videos were acquired using a 4X objective with 4 by 4 camera binning, corresponding to a calibrated image scale of 6.5 µm/pixel. Bright-field recordings were analyzed using custom Python software by tracking the distance between the flexible PDMS posts throughout the contraction cycle. The relaxed post-to-post distance was determined at the diastolic state, and the maximum change in distance was measured at peak contraction. Post deflection was then converted to contractile force using beam bending mechanics, as described previously [25]. The PDMS posts used to form µHT were 1 mm in height with a 270 µm base diameter and a -2° taper, with posts-to-post distance of 1850 µm and had an elastic modulus of 3000 kPa and calculated bending stiffness of 1.96 N/m, corresponding to approximately 98 µN of applied force at 50 µm deflection [25]. Post bending stiffness was characterized based on post geometry and finite-element modeling. Peak active force (µN) was calculated from maximum contraction induced changes in post-to-post distance using:

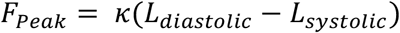

Where *F_Peak_* is peak active force (µN), *κ* is the post bending stiffness (1.96 N/m), *L_diastolic_* is the post-to-post distance at rest (µm), and *L_systolic_* is the post-to-post distance at peak contraction (µm). To account for differences in tissue geometry, we measured the cross-sectional area of each μHT (**Fig 6**) and calculated active stress (**Fig 6**) according to:

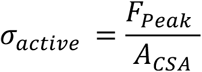

Where *σ_active_* is peak active stress (mN/mm^2^), *F_Peak_* is peak active force (μN), and *A_CSA_* is tissue cross-sectional area (mm^2^). Representative calcium transient and force traces were plotted for each condition.

### Myofibril alignment and contraction kinetics analysis

Myofibril alignment was quantified as the orientational order parameter (OOP), which ranges from 0 for random orientation to 1 for perfect alignment [67, 68]. OOP was calculated from the local orientation of traced myofibril centerlines, measured by structure tensor analysis and pooled across all analyzed planes.

For each beat, contraction velocity was taken as the maximum rate of change of estimated fractional shortening of three randomly selected ROI within each cardioids, between the beat’s onset and its peak, averaged across beats. Estimated circle fitted to the gel boundaries and rate of the changes of estimated circle radios was evaluated.

### Statistical analysis

All statistical analyses were performed using GraphPad Prism (version 10). For every experiment, we distinguish N, the number of independent biological replicates (independent hiPSC encapsulation and differentiation batches), from n, the number of technical replicates (individual cells, gels, colonies, regions of interest, or tissues) measured within those batches; both are reported in each figure legend. Data are presented as mean ± SD for normally distributed data or median with IQR for non-normally distributed data, as indicated in figure legends. Normality was assessed using the Shapiro-Wilk test. Outliers were identified and removed independently for each morphological parameter (cell area, elongation, and circularity) using the ROUT method (Q = 1%) prior to statistical analysis, resulting in slightly different final sample sizes across measurements. For comparisons among three or more groups, one-way ANOVA with Tukey’s post-hoc test was used for parametric data, and Kruskal-Wallis test with Dunn’s multiple comparisons post-hoc test was used for non-parametric data, when each data point represented an independent biological replicate. For two-group comparisons, unpaired t-test with Welch’s correction was used for parametric data, and Mann-Whitney test was used for non-parametric data, under the same condition. Where technical replicates were pooled across multiple independent biological replicates (batches), we instead used nested statistical tests, nested t-test for two-group comparisons or nested one-way ANOVA with Tukey’s or Dunn’s multiple comparisons post-hoc test for three or more groups, to account for this hierarchical data structure and avoid pseudoreplication. Given the large number of cells in each µHT and the independent formation of each tissue, even from the same differentiation batch, we analyzed functional µHT functional data using standard, non-nested one-way ANOVA or Kruskal-Wallis tests with Dunn’s multiple comparisons. Statistical significance was set at α = 0.05, with significance levels indicated as *p < 0.05, **p < 0.01, ***p < 0.001, and ****p < 0.0001.

## Acknowledgments

This work was performed with support from the National Institutes of Health (R21AR082992, R01HL159094 and R01HL107594 to NH, T32 EB028092 to MH) and the National Science Foundation (CAREER 2338931 to NH).

## Author contributions

Conceptualization: MH; Methodology: MH, NH; Investigation: MH, NDD, YKGK, CC; Formal analysis: MH, YKGK, CC, BB, JV; Software: GM, BB; Writing original draft: MH, NH; Writing review & editing: MH, NH; Supervision and funding acquisition: NH.

## Data Availability Statement

All data supporting the findings of this study are available within the main text and supplementary materials. Custom analysis scripts used for morphological quantification are available at https://github.com/huebschlab and mirrored on the lab website (huebschlab.wustl.edu).

