## Supplementary Information for "Xeno-Free Peptide-Functionalized Hydrogels Support hiPSC Encapsulation and *In Situ* Differentiation into Cardiomyocytes with Enhanced Structural Maturation"

**Supplementary Table 1. Key resources table**

| Antibodies and markers and dyes | Source | Identifier |
|---|---|---|
| Sarcomeric-αactinin (clone EA-53) | Sigma-Aldrich | A7811 |
| Sarcomeric-αactinin (polyclonal) | Abcam | AB137346 |
| cardiac troponin T (clone 13-11) | Invitrogen | MA5-12960 |
| Nanog (polyclonal) | Cell Signaling | 35805 |
| MYH7 (clone A4.840) | Santa Cruz Biotech | sc-53089 |
| Desmin (clone D93F5) | Cell Signaling | 5332 |
| Alexa Fluor 488, 568, and 647 conjugated goat anti-mouse secondary antibodies | Invitrogen | A-11001, A-11004, A21235 |
| Hoechst 3342 | Invitrogen | 14533 |
| Zombie dye | Biolegend | 423113 |
| Alamar Blue | Fisher Scientific | Y00-100 |
| LIVE/DEAD® viability kit | Invitrogen | L3224 |

**Supplementary Table 2. Cardiomyocyte yield per input hiPSC across established 2D and 3D hiPSC-CM differentiation platforms.**

| Study | Platform | Pre-diff. seeding day | CM purity (cTnT/TNNT2+) | CMs per input hiPSC | Notes |
|---|---|---|---|---|---|
| **Lian et al., PNAS 2012 (GiWi protocol)** [1] | 2D monolayer | Day −5 | up to 98% | ~15 | Seeded 1–2×$10^5$ cells/cm$^2$. |
| **Burridge et al., Nat Methods 2014 (CDM3)** [2] | 2D monolayer | Day −4 | 85% raw; >95% after metabolic (lactate) selection | ~44 avg.; up to 100 | Seeded 1.25×$10^4$ cells/cm$^2$, reaching ~85% confluence at Day 0; explicitly reports “up to 100 CMs per input pluripotent cell.” |
| **Hamad et al., Theranostics 2019** [3] | 2D monolayer | Day −4 | 91.2% | not stated as a ratio (0.42×$10^6$ CM/cm$^2$) | Same study/protocol also run in suspension (row below). |
| | 3D suspension bioreactor | Day −4 | 87.2% | ≈2.3 (calculated) | 125 mL culture inoculated with 40×$10^6$ hiPSCs → 71.9±29.8×$10^6$ CMs per 100 mL harvested. Same pre-diff. window as their own 2D row. |
| **Kempf et al., Stem Cell Reports 2014**[4] | 3D suspension (stirred bioreactor) | Day −4 | 53.9–84.1% | not reported | Inoculated at 3.3×$10^5$ cells/mL; ~40–50×$10^6$ CMs per 100 mL batch. No per-input-cell ratio given. |
| **Fonoudi et al., Stem Cells Transl. Med. 2015**[5] | 3D suspension (spinner flask) | Day −5 (tested 3/5/7 d; 5 d optimal) | ~90% | not reported | Seeded 2×$10^5$ cells/mL as spheroids; no total-yield or per-input-cell figure given. |

| | | | | | |
|---|---|---|---|---|---|
| **Halloin et al., Stem Cell Reports 2019**[6] | 3D suspension (chemically defined) | Day −2 (optimal; Day −4 tested — underperformed) | 93.5% (150 mL scale); 89–92% (350–500 mL scale) | not reported | In this study the shorter preculture (Day −2 ) outperformed Day −4. |
| **Prondzynski et al., Nat Commun 2024** [7] | 3D suspension (stirred bioreactor, 100 mL) | ~Day −1 (EB-size dependent) | ~94% | 2.4 | Averaged across 25 differentiations, 14 hiPSC lines |
| | 3D suspension (spinner flask, 100–380 mL) | | 94±1.8% | 3.6 (100 mL) → 6.9 (380 mL) | Ratio varies with vessel scale, higher at larger volumes |
| **Hashemi et al., Bioact. Mater. 2025 (first author's prior work)** [8] | 3D PEG-fibrinogen hydrogel microspheres | Day −3 | 68.4 ± 17.7% (Un-Arc 16 FacsII); 71.9 ± 5.5% (IMR90) | 2.1 ± 1.8 (Un-Arc 16 FacsII); 0.9 ± 0.02 (IMR90) | Same Day −3 pre-differentiation seeding window as this study |
| **Hashemi et al. (this study)** | 3D alginate hydrogel (cRGD-functionalized) | Day −3 | ~72% | ~12 avg | Day −3 pre-differentiation seeding (this study) |

**Supplementary Table 3. Consistency of cardiomyocyte differentiation for a single user: 60 kDa 3D EiC vs. optimized 2D differentiation protocol.**

| Platform | Differentiations attempted | Differentiations yielding beating cells | Success rate (%) |
|---|---|---|---|
| 60 kDa 3D EiC | 6 | 6 | 100% |
| 2D (TCPS, optimized protocol) | 6 | 3 | 50% |

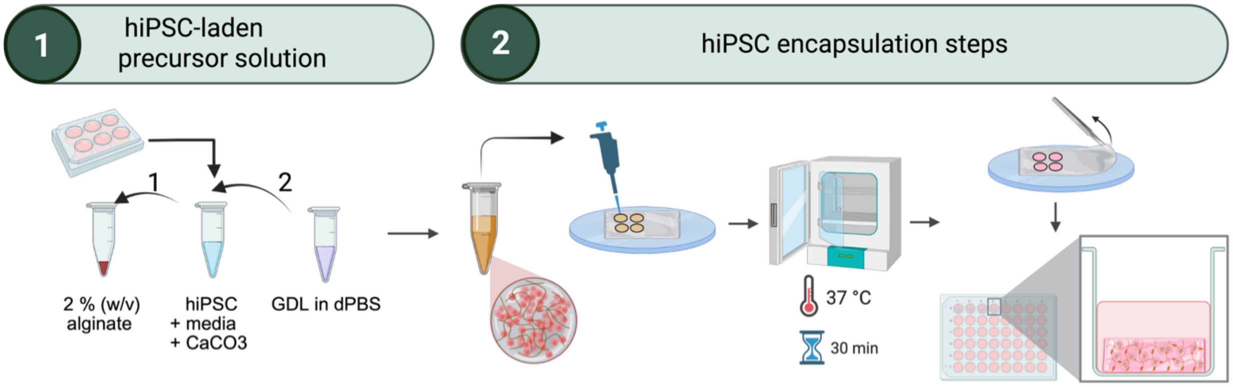


**Supplementary Figure 1. Schematic of the hiPSC encapsulation workflow.** Preparation of the hiPSC-laden precursor solution. A single-cell suspension of hiPSCs in media is combined with $CaCO_3$ and 2% (w/v) alginate to form the cell-laden precursor; GDL in dPBS is prepared separately as the acidifying agent. The precursor solution is mixed with GDL to initiate internal gelation, pipetted as droplets onto a substrate, and crosslinked at 37 °C for 30 min. The resulting hiPSC-laden alginate gels are transferred to culture plates for subsequent expansion/differentiation. Schematics are created with BioRender.

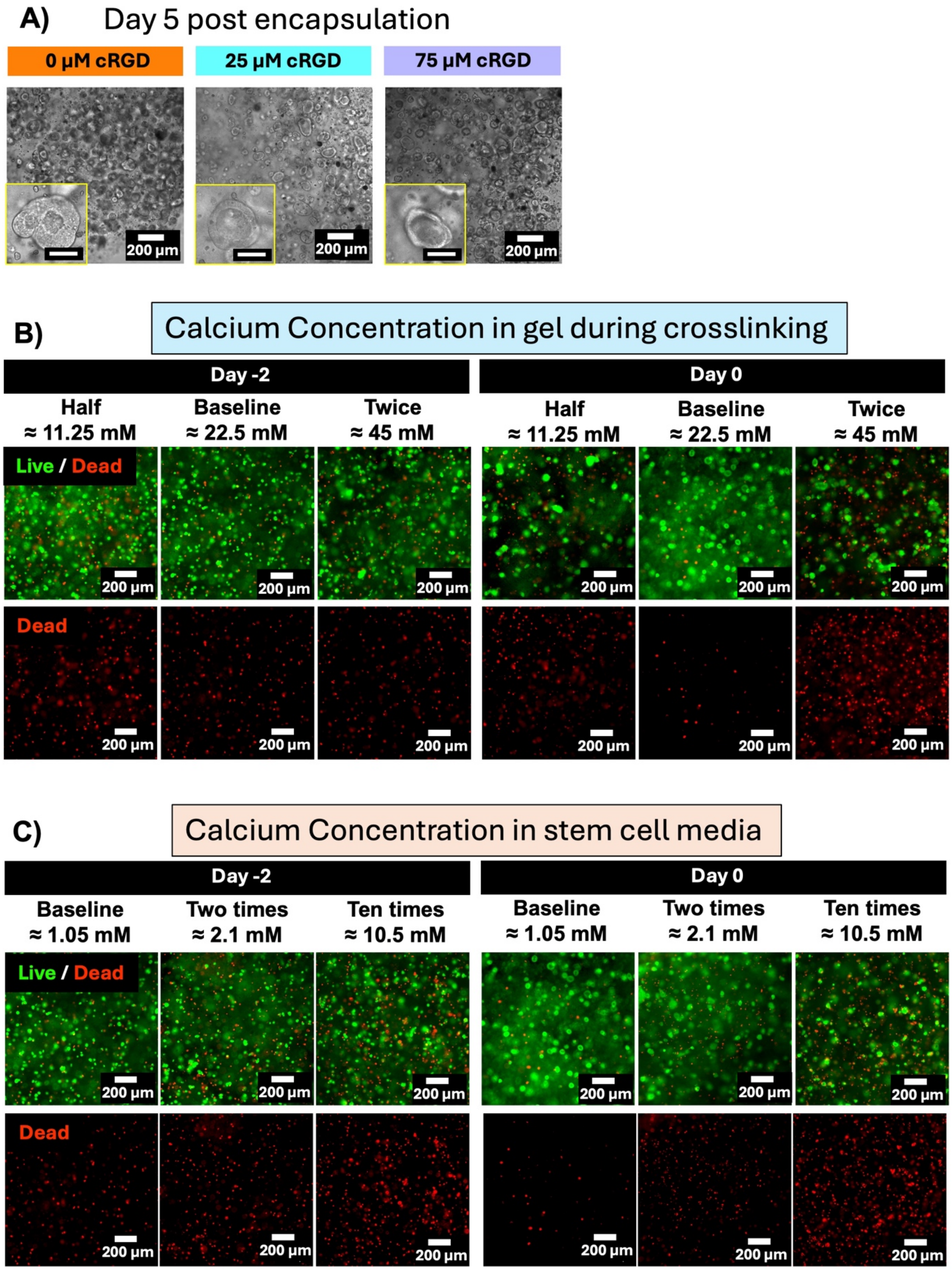


**Supplementary Figure 2. hiPSC viability and morphology across cRGD concentration, calcium crosslinking concentration, and calcium concentration in culture media.** A)

Representative brightfield images of hiPSCs encapsulated in 290 kDa alginate gels functionalized with 0, 25, or 75 µM cRGD, imaged at day 5 post-encapsulation. Insets show magnified single colonies. Scale bars = 200 µm (main) and 50 µm (insets). **B)** Representative Live/Dead (top) and Dead-only (bottom) images of hiPSCs encapsulated in alginate gels crosslinked with half (approximately 11.25 mM), baseline (approximately 22.5 mM), or twice (approximately 45 mM) the standard calcium concentration, imaged at day −2 and day 0. Scale bars = 200 µm. **C)** Representative Live/Dead (top) and Dead-only (bottom) images of hiPSCs cultured in stem cell media containing baseline (approximately 1.05 mM), two times (approximately 2.1 mM), or ten times (approximately 10.5 mM) the standard calcium concentration, imaged at day −2 and day 0. Scale bars = 200 µm.

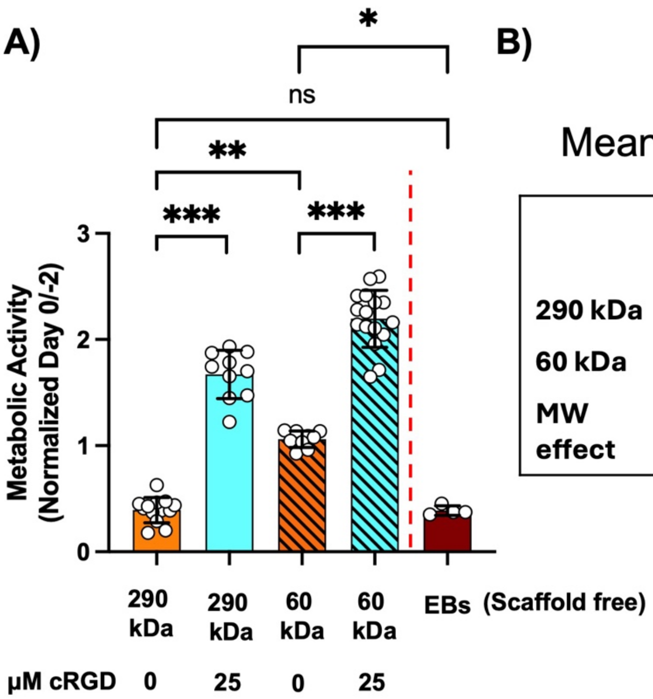


B)

Mean metabolic activity for gels

| | 0 µM cRGD | 25 µM cRGD | cRGD effect |
|---|---|---|---|
| **290 kDa** | 0.39 | 1.67 | +1.28 (~4.3×) |
| **60 kDa** | 1.06 | 2.19 | +1.13(~2.1×) |
| **MW effect** | +0.67 (~2.7×) | +0.52 (~1.3×) | |

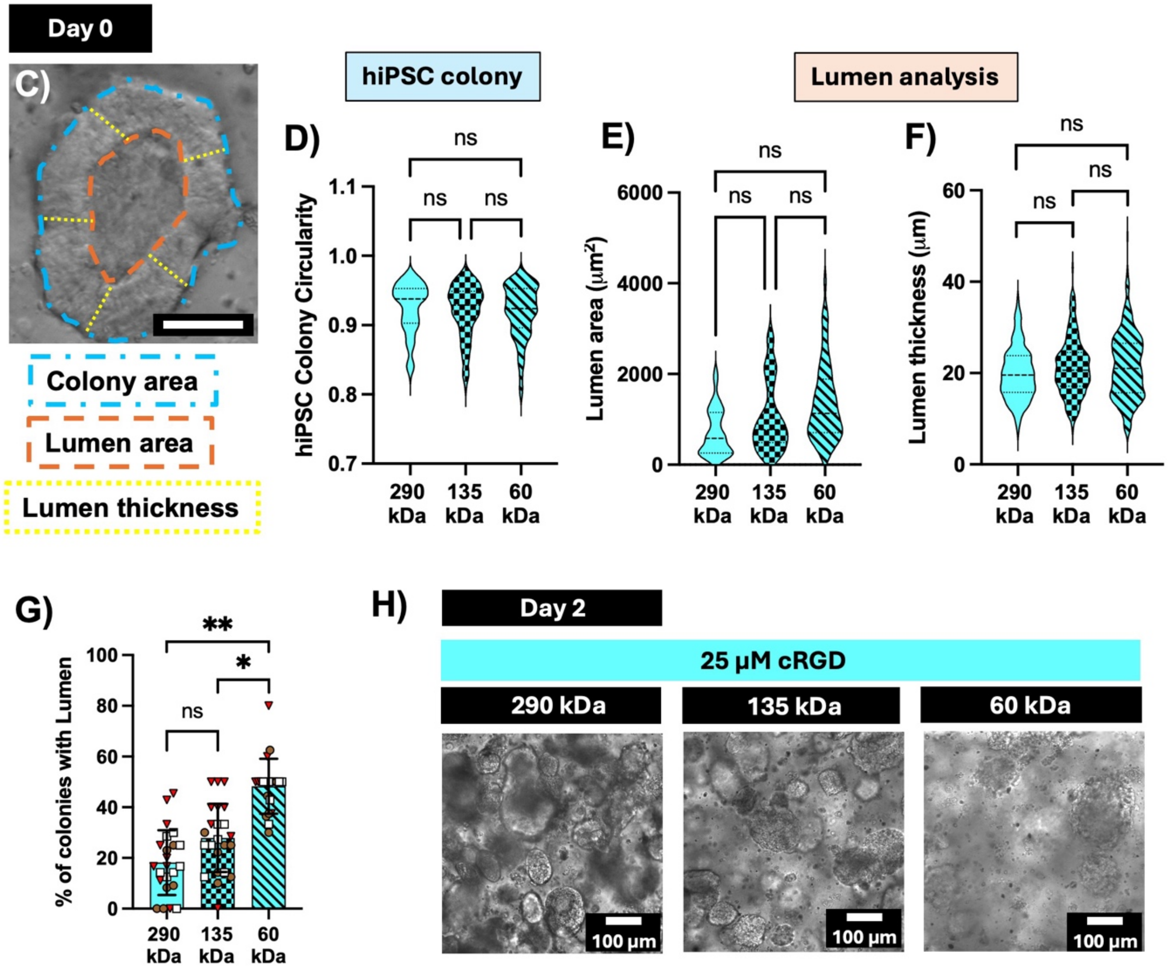

**Supplementary Figure 3. Metabolic activity of encapsulated hiPSCs across gel molecular weight and cRGD concentration, and lumen formation at day 0. A)** Metabolic activity at day 0, normalized to day −2, for hiPSCs encapsulated in 290 kDa (0 and 25 µM cRGD) and 60 kDa (0 and 25 µM cRGD) alginate gels, and in Embryoid Bodies (EBs, scaffold-free). Bars represent mean ± SD. Statistical comparisons were performed using a nested one-way ANOVA with Tukey's multiple comparisons test, treating gels as nested within three independent batches for the scaffold-based conditions. **B)** Mean metabolic activity for 290 kDa and 60 kDa gels at 0 and 25 µM cRGD, with the calculated cRGD effect (addition of cRGD) and MW effect (reduction in molecular weight) on metabolic activity, expressed as absolute difference and fold change. **C)** Representative day 0 brightfield image at 25 µM cRGD illustrating the quantification scheme used in D through G: colony area (blue dashed outline), lumen area (orange dashed outline), and lumen thickness (yellow dashed outline). Scale bar is 50 µm. **D)** hiPSC colony circularity, **E)** lumen area, and **F)** lumen thickness at day 0, at 25 µM cRGD, across three alginate molecular weights (290, 135, and 60 kDa); nested one-way ANOVA, N = 3 independent batches per condition; all comparisons ns. **G)** Percentage of colonies with a lumen at day 0, at 25 µM cRGD, across three alginate molecular weights; nested one-way ANOVA with Tukey's multiple comparisons test, N = 3 independent batches per condition; 290 vs. 60 kDa and 135 kDa vs. 60 kDa significant, other comparisons ns. **H)** Representative brightfield images at day 2 across three alginate molecular weights (290, 135, and 60 kDa) at 25 µM cRGD. Scale bars = 100 µm.

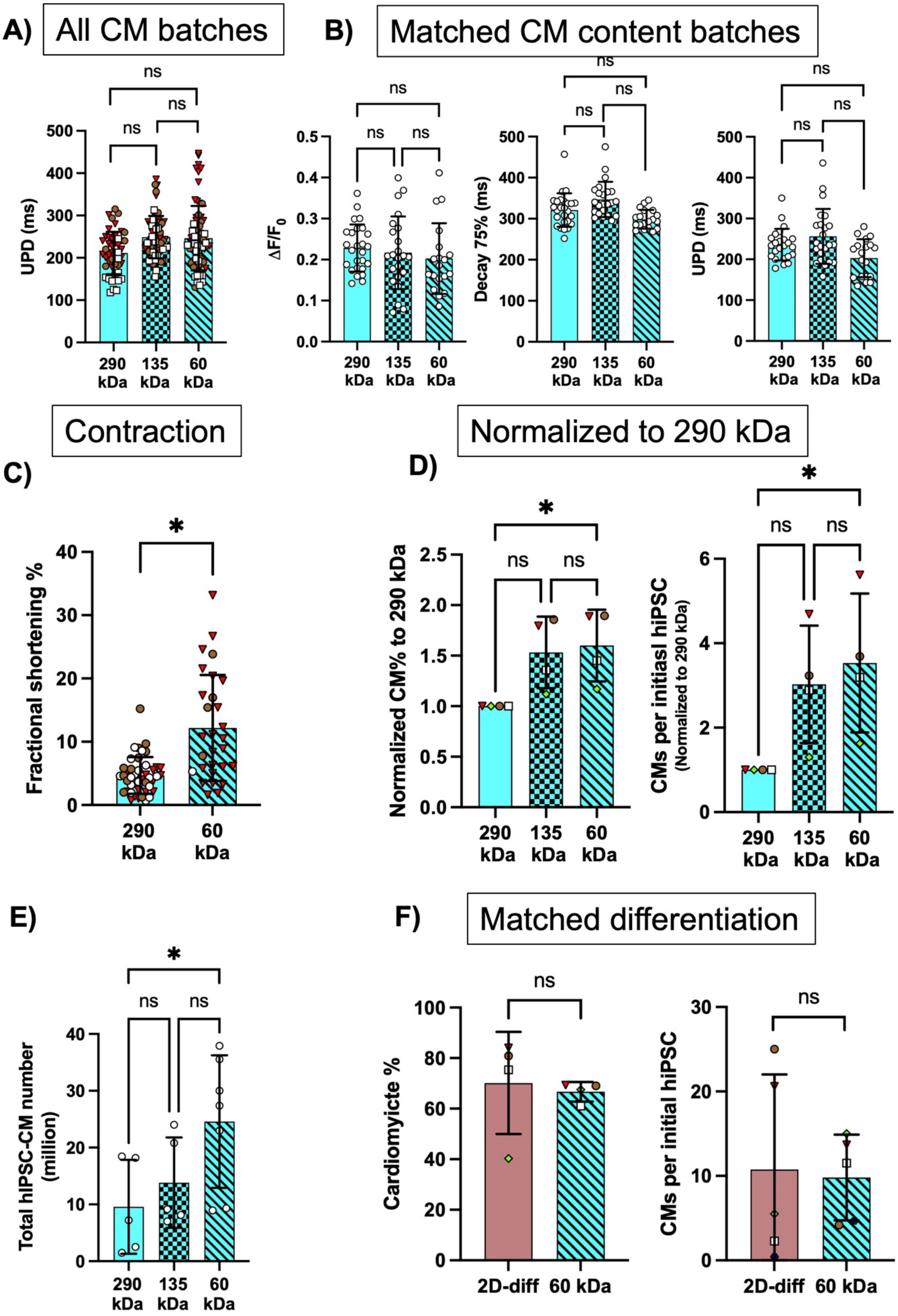
A)
All CM batches
B)
Matched CM content batches
UPD (ms)
ΔF/F0
Decay 75% (ms)
290 kDa
135 kDa
60 kDa
ns
Contraction
C)
Fractional shortening %
*
Normalized to 290 kDa
D)
Normalized CM% to 290 kDa
CMs per initiasl hiPSC
(Normalized to 290 kDa)
E)
Total hiPSC-CM number (million)
F)
Matched differentiation
Cardiomyicte %
CMs per initial hiPSC
2D-diff
60 kDa

**Supplementary Figure 4. Functional and yield comparison of hiPSC-derived cardiomyocytes across gel molecular weights. (A)** Upstroke duration (UPD) of EiCs created with 290, 135, or 60 kDa alginates, across all CM batches. **(B)** Functional properties measured in batches matched for cardiomyocyte content: calcium transient amplitude ($\Delta F/F_0$), 75% decay time (ms), and upstroke duration (UPD, ms) across the three gel molecular weights ($n \geq 53$ ROI recordings). **(C)** Fractional shortening (%) of EiCs derived from 290 kDa and 60 kDa alginate gels. **(D)** Cardiomyocyte yield normalized to the 290 kDa condition: cardiomyocyte percentage (Normalized CM%) and cardiomyocytes per input hiPSC. Bars represent mean ± SD ($N \geq 4$ batches). **(E)** Total number of hiPSC-CMs post dissociations. Bars represent mean ± SD ($N \geq 5$ batches). **(F)** Differentiation efficiency in batches matched for differentiation: cardiomyocyte percentage and cardiomyocytes per input hiPSC for 2D-differentiated (2D-diff) and 60 kDa 3D-encapsulated conditions.

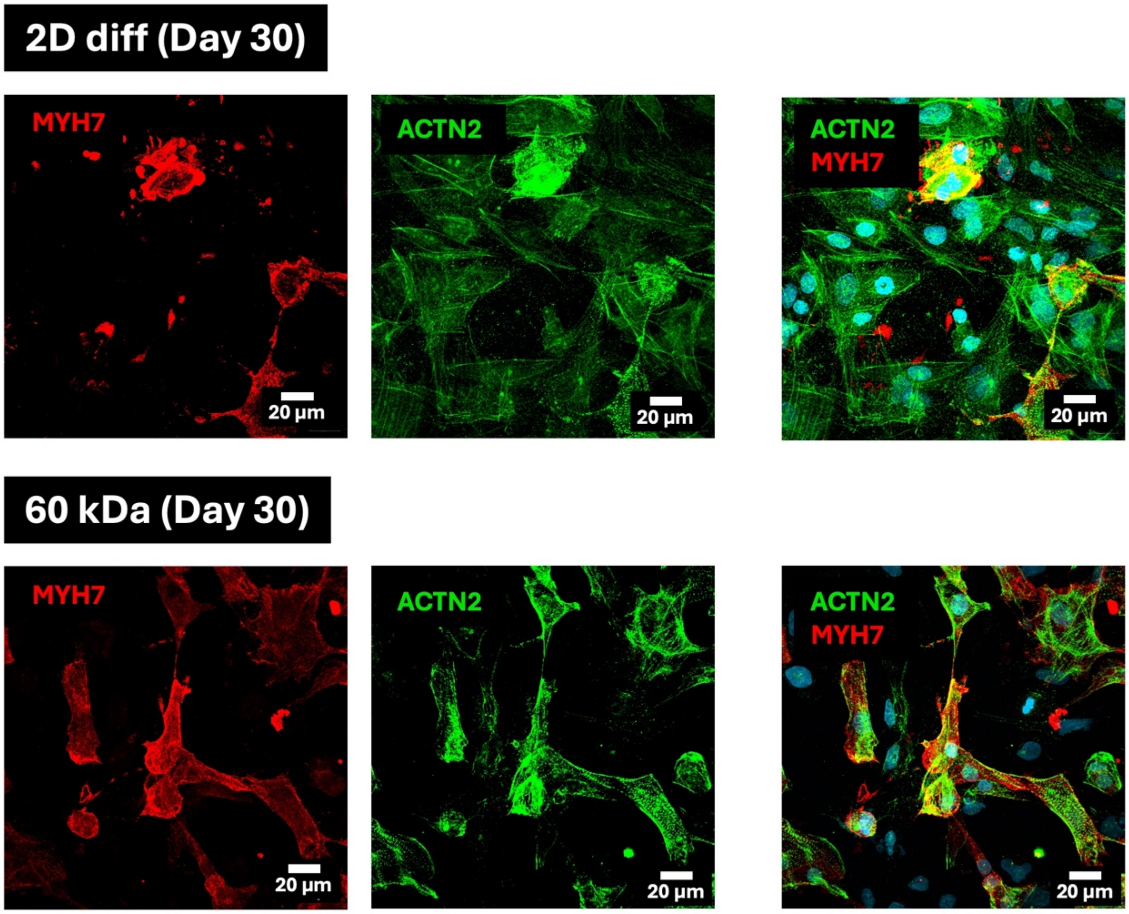


**Supplementary Figure 5.** MYH7 and ACTN2 expression in 2D monolayer- and 60 kDa-derived hiPSC-CMs at day 30. Representative immunofluorescence images of hiPSC-CMs from 2D monolayer differentiation (top) and 60 kDa EiCs (bottom) at day 30, stained for MYH7 (red) and ACTN2 (green), shown separately and merged with nuclei (cyan). Scale bars = 20 µm.

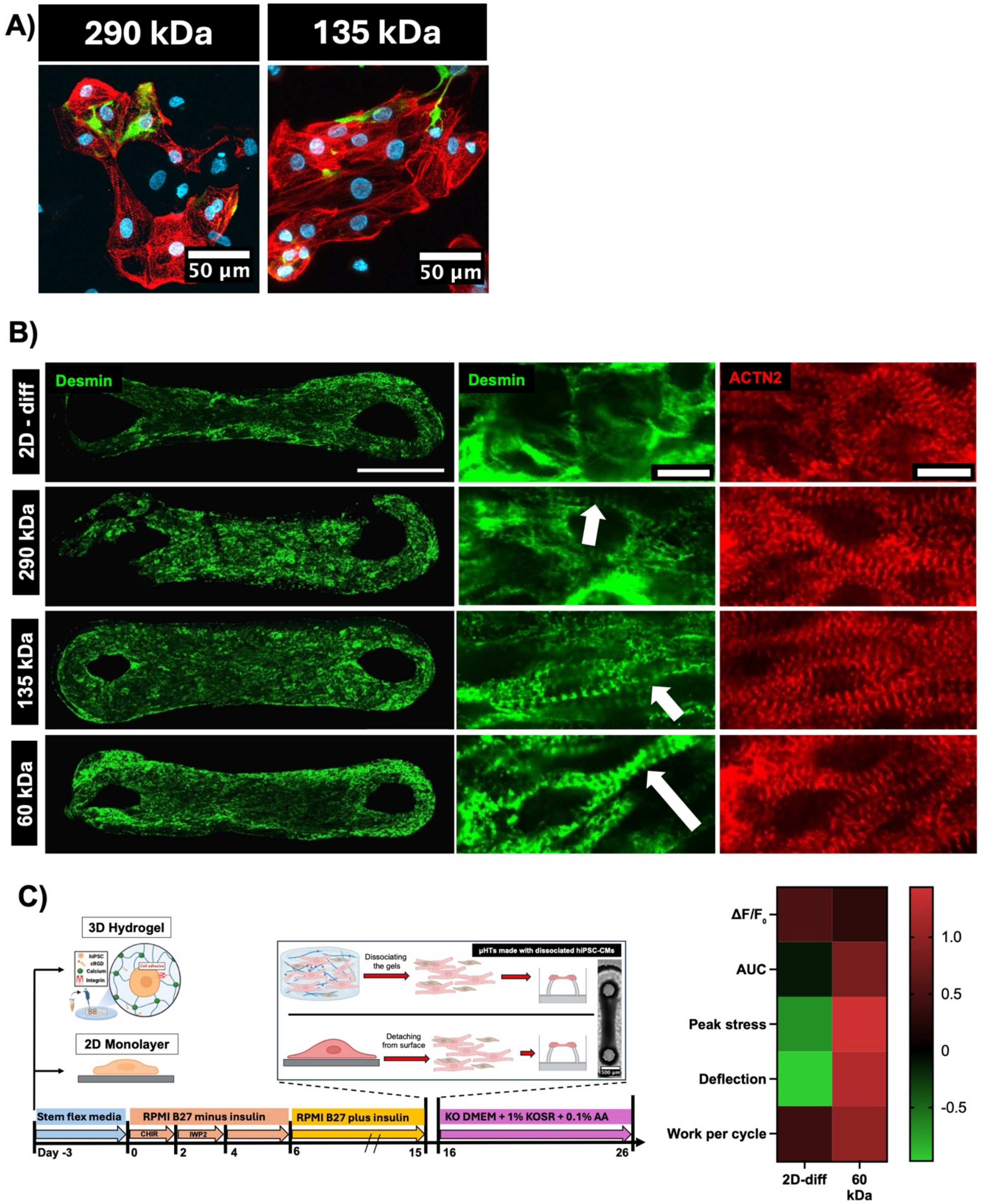


**Supplementary Figure 6.**
**Desmin organization and µHT function of hiPSC-CMs from 2D monolayer and alginate EiC differentiations.**
**A)** Representative immunofluorescence images of dissociated hiPSC-CMs at day 15 stained for Desmin (green), ACTN2 (red), and nuclei (cyan) across 290 kDa and 135 kDa at 25 µM cRGD.

**(B)** Representative immunofluorescence images of Desmin (green) and ACTN2 (red) within intact µHTs fabricated from 2D monolayer-, 290 kDa-, 135 kDa-, and 60 kDa-derived hiPSC-CMs. Insets show higher-magnification views of Desmin and ACTN2; arrows highlight striated Desmin patterning, more prominent in gel-derived conditions than in 2D. Scale bars = 500 µm (whole µHT) and 10 µm (zoomed insets). **(C)** Left, schematic of the 3D hydrogel and 2D monolayer differentiation workflows and the timeline of media conditions (day −3 to day 26); center, generation of micro-heart tissues (µHTs) from dissociated hiPSC-derived cardiomyocytes; right, heatmap summarizing functional readouts ($\Delta F/F_0$, AUC, peak active force, deflection, work per cycle) for 2D-diff and 60 kDa conditions.

**Supplementary Video 1- GCaMP calcium imaging of EiCs at day 15, 290 kDa**

**Supplementary Video 2- GCaMP calcium imaging of EiCs at day 15, 135 kDa**

**Supplementary Video 3- GCaMP calcium imaging of EiCs at day 15, 60kDa**

**Supplementary Video 4_ calcium-wave fluorescence uHT from 290 kDa**

**Supplementary Video 5_ calcium-wave fluorescence uHT from 135 kDa**

**Supplementary Video 6_ calcium-wave fluorescence uHT from 60 kDa**

**Supplementary Video 7_ bright-field uHT from 290 kDa**

**Supplementary Video 8_ bright-field uHT from 135 kDa**

**Supplementary Video 9_ bright-field uHT from 60 kDa**

**Supplementary Video 10_ whole tissue staining 60 kDa**
